# Gene conversion facilitates rapid evolution of inversions across avian immunoglobulin loci

**DOI:** 10.64898/2026.09.05.749481

**Authors:** Katalin Voss, Daniel Hardesty, Anton Zamyatin, Mariia Pospelova, Yixin Zhu, Maria Recuerda Carrasco, Anton Bankevich, Leonardo Campagna, Yana Safonova, Matt Pennell

## Abstract

The genomic architecture of immunoglobulin (IG) loci in birds has received remarkably little attention, despite their relevance to infectious disease susceptibility. One of the few exceptions is the domestic chicken, which has been found to use a completely different mechanism to generate a diverse IG repertoire than most vertebrates; rather than relying on V(D)J recombination, chickens primarily use somatic gene conversion. Whether this is true of all birds has remained unknown and untestable at scale until now. And importantly, it is not known how this alternative mechanism for antibody generation shapes, and is shaped by, genome evolution in birds. Leveraging IG locus annotations from 122 bird species generated through the Vertebrate Genomes Project and 17 species from the California Conservation Genomics Project, we show that avian IGH loci display a striking, previously unreported architecture of recurrent inverted duplications that generate direct and inverted copies of the same repeat unit, found in no other vertebrate lineage. Inversion density varies considerably across species, and population-level analyses reveal that these inversions evolve rapidly. We propose a model in which these inversions are actively maintained because they continuously replenish a pool of highly similar pseudogenes that serve as donors for somatic gene conversion, substituting for the large functional V gene repertoires other vertebrates use to generate IG diversity. This model makes a direct prediction: IGH loci should harbor few functional genes and many pseudogenes, while IGL loci, which typically lack this inversion architecture, should show the opposite pattern. Our cross-species analysis confirms this. To test the model at the level of the expressed repertoire, we generated paired whole-genome and Iso-seq data from a single wild-caught Red-winged Blackbird. Consistent with our predictions, a single terminal IGLV gene is diversified through gene conversion from surrounding pseudogenes, while IGH carries a large donor pool at which we also detect gene conversion. Together, these findings reveal that the molecular evolution of IG in birds is governed by fundamentally different constraints and processes than in the rest of known vertebrates.

## Introduction

B cells and their associated immunoglobulins (IGs) are central components of the adaptive immune system, functioning to bind and neutralize a wide range of pathogens. In most tetrapods, IGs consist of two heavy chains and two light chains, encoded by the heavy chain locus (IGH) and the light chain loci (lambda chain: IGL or kappa chain: IGK), respectively ^1,2^ (however, there is variation among lineages in the number of chains ^3^; see below) The generation of a diverse IG repertoire is essential for the recognition of the broad antigenic spectrum presented by pathogens. In mammals, this diversity is achieved through V(D)J recombination in maturing B-cells, a somatic process by which one variable (V), one diversity (D), and one joining (J) gene are selected from their respective genomic clusters and joined to form the variable region of the heavy chain, with an analogous VJ recombination occurring at the light chain loci ^1,2^. Every functional IG gene requires a recombination signal sequence (RSS) to undergo this V(D)J recombination process to guide the RAG1/RAG2 enzyme complex, which facilitates V(D)J recombination ^2,4^. Studies across mammals have established that these sequences are highly conserved ^5^: each RSS consists of a heptamer located directly downstream of the gene, followed by a 23 bp spacer and a nonamer ^2,4^. Both the heptamer and nonamer follow consistent motifs across mammalian species, with little variation between species ^3,5^. These motifs are so conserved, that they also allow for the discovery of IG genes in non-avian reptiles ^3^.

Due to their highly repetitive and structurally complex nature, the IG loci have historically posed significant challenges for genome assembly ^6–13^. As a result, in-depth analyses have largely been restricted to a small number of model organisms ^14–18^. Bird IG loci in particular have received little attention beyond the domestic chicken (*Gallus gallus*) in stark contrast to other genomic regions involved in the adaptive immune system, most notably MHC, which have been finely characterized over a wide variety of species ^19–23^. Extensive molecular characterization of the chicken IG loci has revealed several unusual features relative to mammals: Where mammals rely on a large repertoire of functional V, D, and J genes to generate IG diversity through combinatorial recombination, the chicken appears to have discarded this strategy almost entirely. Chickens possess only lambda light chains, lacking an IGK locus ^24,25^, and harbor just a single functional V gene and a single functional J gene in both IGH and IGL ^26–29^. Their IGH D gene cluster is notably more homogeneous than that of primates or rodents ^27^ and the RSS associated with the single functional IGH V gene (IGHV) deviates from canonical mammalian motifs ^5,30–32^. Upstream of the single functional V gene in both IGH and IGL loci lies a cluster of pseudogenes arranged in alternating directions ^28,33,34^. Despite this highly restricted germline repertoire, chickens are capable of generating diverse IGs and mounting effective adaptive immune responses. This has been shown to be largely achieved through somatic gene conversion, whereby sequences from upstream pseudogenes are incorporated into already-rearranged V(D)J sequences, diversifying the IG repertoire post-recombination ^33–36^. This process, which occurs in the bursa of Fabricius (a bird-specific lymphoid organ central to avian B cell development ^27,36^), transforms a cluster of non-functional pseudogenes into the main driver of chicken immune diversity. Somatic gene conversion in IG loci is mediated by activation-induced cytidine deaminase (AID) and Rad51 paralogs ^36–40^. AID also drives somatic hypermutation, and the two processes appear to represent competing diversification pathways: when gene conversion is disrupted, either through the deletion of pseudogenes or the loss of Rad51 paralogs, somatic hypermutation increases significantly ^36,37,41,42^. Gene conversion has also been inferred from the preferential use of a single germline V gene in several mammalian lineages^43–46^, though these observations are from short-read data with incomplete germline gene sets and have yet to be confirmed with long-read sequencing methods.

Whether the features of the chicken immune system represent a lineage-specific solution or reflect a broader avian strategy for generating IG diversity is not well-established. Gene conversion has been proposed as a diversification mechanism in a small number of other avian species based on indirect evidence: several non-galliform species, showed a single major IgL rearrangement event, as in the chicken ^47^ and restricted V and J gene usage in the duck has similarly been attributed to gene conversion ^48^. However, this evidence is indirect and based on limited data, leaving open the question of how general this strategy truly is across birds and what it would mean for the molecular evolution of the genes involved. The importance of addressing these questions is underscored by the rise of highly pathogenic avian influenza, which has infected multiple bird species ^49–51^.

As part of Phase I of the Vertebrate Genomes Project ^3^, the IG loci of over 150 birds were annotated and analyzed for the first time, of which 122 possessed high-quality annotations enabling systematic cross-species comparison. We also analyzed an additional 17 species from the California Conservation Genomics Project ^52^. In addition to analyzing these genomes in greater depth, we characterized IG locus variation at the population level using multiple haplotypes from each of nine species. We also collected new paired (i.e. from the same individual) whole-genome sequencing (WGS) with B cell Iso-seq data to directly connect germline gene content with the expressed IG repertoire (from a wild caught Red-winged Blackbird (*Agelaius phoeniceus*)). Using this comprehensive, multi-scale dataset we show that inverted duplications are a hallmark of avian IGH loci across the bird phylogeny, that these inversions are generated rapidly and vary even within species, and that they appear to be actively maintained because they continuously replenish the pseudogene pool required for somatic gene conversion. Together, our findings support a model in which gene conversion, rather than combinatorial V(D)J recombination, generates most IG diversity across birds, with inversions supplying the genomic architecture that makes this possible.

## Results

### Avian IGH have an inversion-driven architecture

The cross-species analysis in Formenti et al. 2026 ^3^ confirmed that the 154 annotated bird species lack an IGK locus as was previously reported in chickens ^24,25^, revealing this is a universal feature of birds rather than a lineage-specific loss. Comparing the IGH locus between mammals, non-avian reptiles, and birds reveals that avian IGH loci are substantially shorter, averaging just 0.2 Mbp compared to 1.7 Mbp in mammals and reptiles (Figure 1A), and this compactness extends to the chromosomal level: avian IGH loci reside on contigs averaging less than 4 Mbp in length, compared to an average of 124 Mbp in mammals and reptiles (Figure S1A). Previous examination of the Pacific Biosciences HiFi coverage for the zebra finch (*Taeniopygia guttata*) IGH locus helps to explain why assembly of these regions is particularly challenging: the locus resides within a homopolymer-rich region enriched with non-canonical (non-B) DNA structures (Supplementary Section 2, Figure S2) that causes a marked drop in HiFi read coverage, likely contributing to local assembly fragmentation ^3^. This difficulty appears specific to IGH rather than a general property of avian IG regions: A systematic analysis of HiFi sequencing depth and sequence composition across 24 species confirmed that IGH, but not IGL, is specifically associated with elevated homopolymer richness and reduced coverage in its chromosomal neighbourhood (see Supplementary Section 2, Figure S2, Table S2.1). To ensure that the assemblies underlying our analysis are of sufficient quality, we restricted our dataset to 122 birds for which both IGH and IGL could be confidently annotated, and applied rigorous filtering at both the contig and gene level to remove likely false positive annotations before any downstream analysis (see Methods).

**Figure 1:**
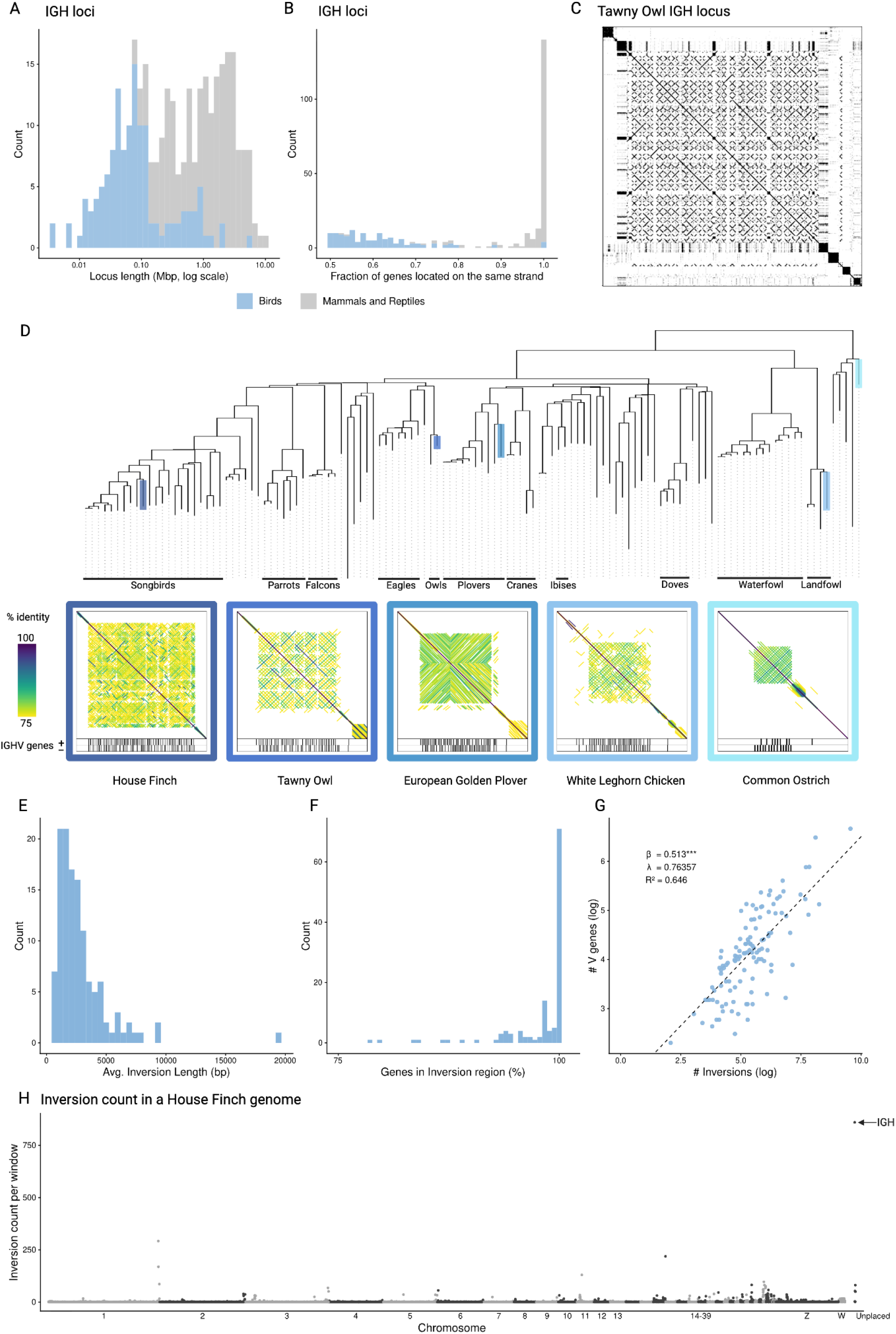
The avian IGH locus A) Locus length comparison between birds (blue) and mammals and reptiles (grey) from the VGP. Length is measured in Mbp and shown on a log scale. B) Fractions of genes located on the same strand compared between birds (blue) and mammals and reptiles (grey) from the VGP. The scale goes from 0.5-1, 1 meaning all genes are located on the same strand and 0.5 meaning half of the genes are on the positive and half on the negative strand. C) Dotplot of a Tawny Owl IGH locus (created with *Gepard* ^81^) D) Species tree of 122 birds from the VGP. Orders with 2 or more species are indicated underneath. Underneath there are 5 example dotplots with the gene positions and strands indicated below for one haplotype of a House Finch (dark blue), Tawny Owl (light blue), European Golden Plover (turquoise), White Leghorn Chicken (dark green) and Common Ostrich (light green). The lines in the dotplots are colored by identity (75%: yellow, 100%: purple). Their position in the tree is indicated in the corresponding colors. E) Histogram of the average inversion length (bp) across all birds F) Histogram of percentage of genes that lie in inversion regions across all birds. G) Relationship between the number of inversions (log) and the number of V genes (log) for all birds. The phylogenetic regression was calculated using *phylolm* ^82,83^ under the lambda model and the relationship was found to be significant (beta=0.513***, lambda=0.76357, R^2=0.646). H) Inversion count per window of the same size as the IGH locus (108.5 kbp) across the genome of a house finch. The dot representing the IGH locus is marked with an arrow.

While the majority of non-avian species show a clear directional preference in IGH gene orientation, birds show a strikingly equal distribution of genes in both orientations (Figure 1B), a pattern reminiscent of the alternating pseudogene arrangement previously described in the chicken and suggesting that this too might be a general and conserved feature across birds. The picture is different for IGL: although the pseudogenes in the chicken were already known to have genes in both directions in IGL ^22,24,25^, our cross-species analysis reveals this to be unusual among birds more broadly, with the vast majority of species retaining a clear direction bias in IGL (Figure S3). There are just two exceptions to this rule: all landfowl (including the chicken) in our dataset show an equal distribution of genes in both directions in IGL, as does a second, distantly related clade (Daedalornithes) that includes the Whiskered Treeswift (*Hemiprocne comata*), Anna’s Hummingbird (*Calypte anna*) and the Mountain Owlet-nightjar (*Aegotheles albertisi*) in our dataset. That two such phylogenetically distant groups independently arrived at the same unusual IGL architecture hints at convergent selective pressures that remain to be understood.

To investigate the genomic basis of this unusual gene orientation pattern, we generated dotplots of the IGH loci for each haplotype across a phylogenetically diverse set of bird species (Figure 1C, Figure 1D). The dotplots reveal that IGH genes alternate in orientation because the locus is riddled with inverted duplications: segments ranging from single genes to clusters of several genes are copied and reinserted in reverse orientation, so that both the original segment and its inverted copy persist, placing the duplicated genes on the opposite strand from the surrounding sequence. A segment and its reverse-complement copy together form a palindrome (an inverted repeat), and because each event both flips and copies the sequence, these inverted duplications progressively expand the locus while generating the alternating-strand, inversion and palindrome-rich pattern we observe. The resulting dotplots show a distinctive, densely crossed pattern resembling cross-stitch, as the accumulation of many inverted duplications of different sizes and positions repeatedly flips the diagonal alignment back and forth across the locus (Figure 1C). This pattern was distributed broadly across the bird phylogeny, indicating that a high frequency of inversions is a common feature of the avian IGH locus. To our knowledge, no analogous inversion architecture has been reported at the IGH locus of any other vertebrate lineage, marking this as a derived feature unique to birds and raising the question of what evolutionary forces have driven and maintained it. The number of inversions varies considerably across species: the house finch, for example, harbors a substantially higher number than the ostrich in the representative haplotypes, as illustrated by the representative dotplots in Figure 1C. Individual inversions are also variable in length, though the majority are shorter than 5 kbp with an average of 2.8 kbp (Figure 1E). Notably, on average 98% of all IGHV genes in a given species lie in the inversion associated region (Figure 1C, 1F), and inversion count and V gene number show a strong positive correlation (β = 0.513***, λ=0.76, R^2^=0.646) (Figure 1G), suggesting that inverted duplications are a primary mechanism through which birds expand their IGHV gene repertoire. Remarkably, despite this variability, the inversion architecture remains recognizably alignable even between species separated by deep divergences (Figure S4), whereas in mammals IGH loci diverge too rapidly to retain comparable cross-species alignment^32^. Avian IGH loci are thus simultaneously highly dynamic and, in their underlying repeat structure, unexpectedly conserved across the phylogeny.

This inversion pattern is largely absent from IGL. IGL loci are also substantially shorter than those of mammals and non-avian reptiles (Figure S5.1A). This compactness in both IG loci is likely a reflection of a more general feature of avian genomes, which are known to be more compact overall and characterized by a greater number of micro and dot chromosomes than other vertebrates ^53–55^, the short chromosomes on which avian IGH loci appear to predominantly reside. Consistent with the strand bias analysis (Figure S3, Figure S5.1B), we do find inversions in IGL in the same clades that lack a strand bias there, namely landfowl and the swifts, hummingbirds and their relatives, confirming that the two observations reflect the same underlying phenomenon. Beyond these exceptions, a genome-wide comparison of inversion frequency across loci of equivalent length in the housefinch reveals that no other locus comes close to the inversion density observed at IGH (Figure 1H), underscoring how unusual and likely functionally significant this architecture is.

### IGH inversion architecture evolves rapidly within species

With the sequence-level contributors to IGH assembly difficulty established, we turn back to the inversions themselves to ask how rapidly they arise and how much they vary within a species. To address this, we took advantage of population-level pangenome datasets for nine species with multiple haplotypes available: House Finch (*Haemorhous mexicanus*, n=33) ^56^, Island Scrub Jay (*Aphelocoma insularis*, n=15), Florida Scrub Jay (*Aphelocoma coerulescens*, n=14), Woodhouse’s Scrub Jay (*Aphelocoma woodhouseii*, n=14)^57^, Ibera Seedeater (*Sporophila iberaensis*, n=8), Marsh Seedeater (*Sporophila palustris*, n=6), Tawny-bellied Seedeater (*Sporophila hypoxantha*, n=6), Chestnut Seedeater (*Sporophila cinnamomea*, n=4), and Dark-throated Seedeater (*Sporophila ruficollis*, n=4)^58^. Aligning the IGH loci within each species revealed a striking range of within-species diversity (Figure 2, Figure S6.2, Figure S6.3). The Island Scrub Jay, an endemic species restricted to Santa Cruz Island off the California coast with a small and isolated population ^57^, shows essentially identical haplotypes (Figure S6.1).

**Figure 2:**
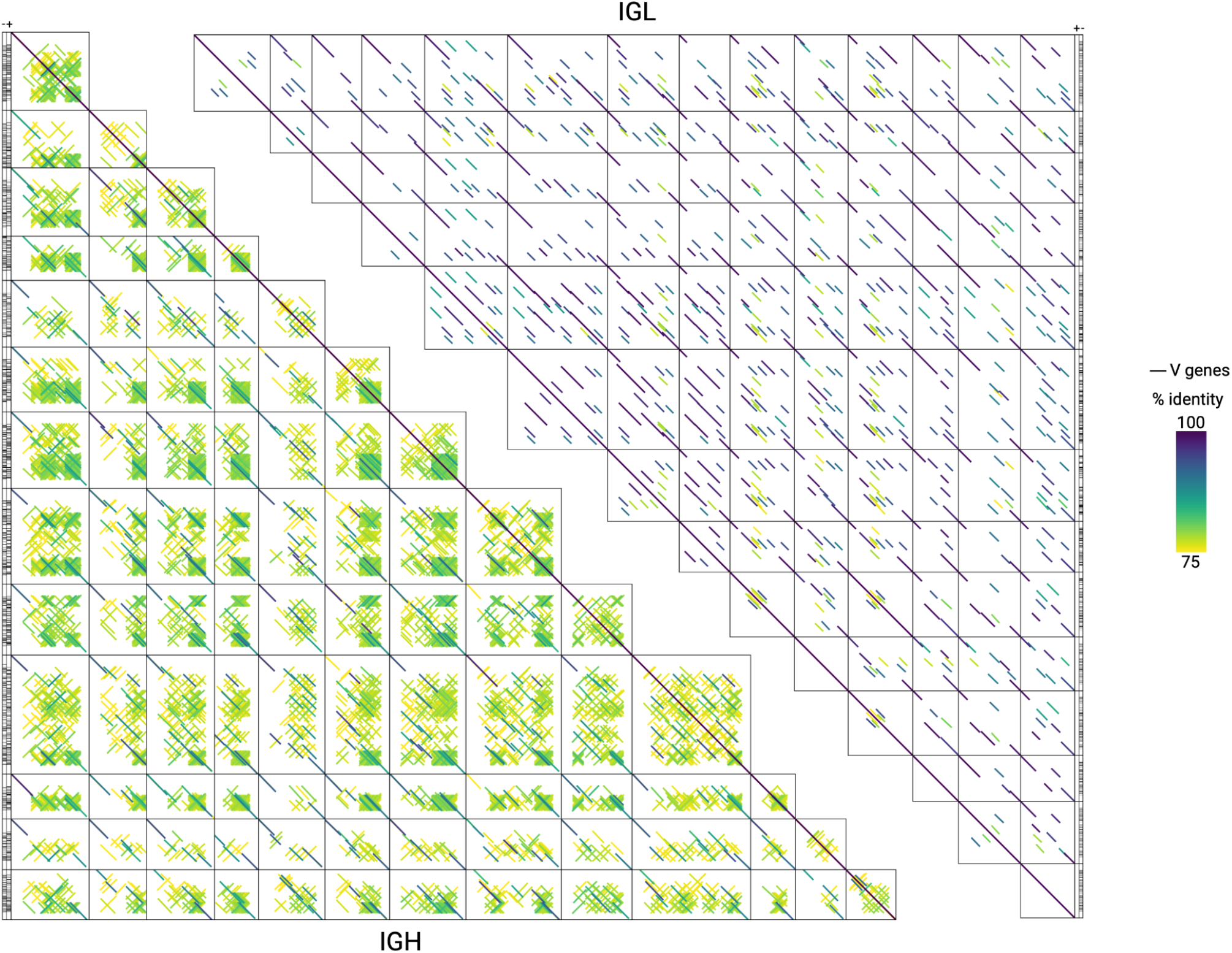
Pairwise whole-locus alignments of the IGH (13 haplotypes) and IGL (14 haplotypes) loci across primary genome assemblies of Woodhouse Scrub-Jay (Aphelocoma woodhouseii) individuals. Each matrix is arranged as a triangle: diagonal panels show a locus aligned against itself, while panels off the diagonal show pairwise alignments between two different individuals’ loci. Every line is a local nucleotide alignment (LASTZ) of at least 10 kb between the two sequences plotted on that panel’s axes, colored by percent identity as indicated in the color bar (75%: yellow, 100%: purple). Each line in a non-diagonal panel marks sequences shared between the two haplotypes. Black bars alongside each locus mark the positions of annotated V genes, with the strand indicated at the top.

This also provides strong evidence that the inversion patterns we observe reflect true biological signal rather than assembly error, as we would not expect to see the same inversion pattern across haplotypes if this was an error. The Woodhouse’s Scrub Jay, by contrast, is widely distributed across western North America and has an effective population size approximately 60 times larger than that of the Island Scrub Jay ^57^. Consistent with this, its IGH loci show considerably more diversity across haplotypes (Figure 2), though shared inversions are still present. Together, these species bracket the range of within-species IGH locus diversity in our dataset.

The same species also lays bare the contrast between the two loci. In the Woodhouse Scrub Jay, IGL is also not static across haplotypes. Full-length diagonal alignments are rare, and the locus instead breaks into many short, often offset segments, indicating substantial within-species variation there too. What IGL lacks is the dense inversion structure that pervades IGH: the heavy chain locus shows the same underlying fragmentation plus extensive turnover of inverted segments layered on top (Figure 2). Both loci diversify within a species, yet only IGH does so through recurrent inversion.

### Chromosomal context helps explain why inversions are restricted to IGH

The observation that inversions are both widespread across species and variable within them raises a deeper question: why have they arisen and been maintained so extensively at IGH locus throughout bird evolution? We propose two contributing factors: IGH’s chromosomal neighborhood makes it unusually prone to being included in inversions, and gene conversion supplies a benefit large enough to offset their cost.

IGH resides on small dot chromosomes whose local sequence composition differs markedly from the rest of the genome: the region surrounding IGH is enriched for low-complexity, homopolymer-rich sequence (Supplementary Section 2, Figure S2), which promotes hairpin and other non-B DNA structures that stall replication ^59–63^. Resolution of these stalls through Fork Stalling and Template Switching (FoSTeS) or Microhomology-Mediated Break-Induced Replication (MMBIR), both associated with short inversion formation ^60^, would repeatedly generate new inverted repeats, offering a plausible explanation for why inversions arise so frequently at this locus, largely independent of downstream selection. Consistent with this, the central portions of palindromic inversions in IGH are less palindromic than their flanking sequences (Supplementary Section 7, Figure S7), suggesting a non-palindromic spacer, as predicted if these structures form through microhomology-mediated template switching.

Inversions are generally destabilizing to genomes ^59,64–66^, yet they are abundant and persistent at IGH, implying that some benefit offsets this cost at this locus. We propose that this benefit is gene conversion. Gene conversion depends on substantial sequence homology between the functional gene and its donor ^34^, a resource the duplicated, inverted copies generated by these events directly supply. Gene conversion in the chicken has further been shown to preferentially utilize donor genes on the opposite strand from the functional gene ^34^, precisely the configuration inversions repeatedly produce. Inversions thus let birds sustain a diverse repertoire from a physically short locus without a large functional V gene family, and we suggest this combination, frequent inversion formation paired with a mechanism that both requires and preferentially draws on the resulting duplicates, is what has allowed inversions to accumulate at IGH rather than being purged as at other loci.

This leaves one observation unexplained: gene conversion operates at both IGH and IGL in the chicken ^33,35,36,42,67–69^, yet the inversion pattern is exclusive to IGH in most birds. Under our model this follows from chromosomal context rather than any difference in the benefit gene conversion provides at the two loci. IGL resides on larger chromosomes (Figure 3A) and, unlike IGH, shows no enrichment for homopolymeric sequence in its local neighbourhood (Supplementary Section 2, Figure S2C): the sequence context that appears to drive inversion formation at IGH is largely absent at IGL. Consistent with the overall pattern, IGL V genes are almost always restricted to a single strand, with the exception of landfowl and Daedalornithes (Figure 3B, Figure S3), and inversions in IGL occur precisely in these same clades, confirming the two observations reflect the same underlying phenomenon. We speculate that the distribution of fitness effects of inversions will likely be different between the micro-chromochromes and the larger chromosomes owing to differences in the number and type of genes and this will contribute to whether a new inversion becomes fixed; however, investigating this is beyond the scope of the present analysis.

**Figure 3:**
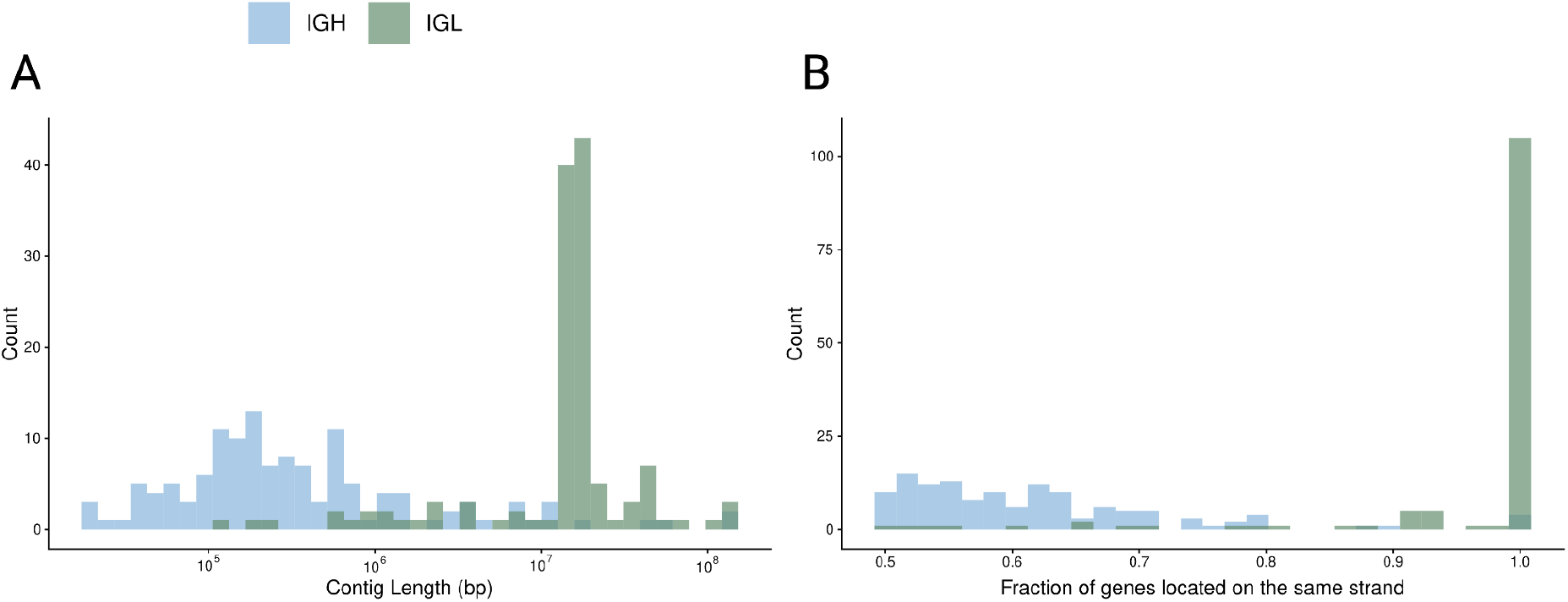
Comparisons between IGH and IGL A) Length (bp) of the contigs that IGH and IGL lie on (log scale) B) Fraction of genes located on the same strand compared between IGH and IGL. The scale goes from 0.5-1, 1 meaning all genes are located on the same strand and 0.5 meaning half of the genes are on the positive and half on the negative strand.

Together, these two effects are likely two pieces of the same puzzle: an ancestral translocation event placing IGH onto a low-complexity local environment on a small dot chromosome. Large-scale chromosomal rearrangements of this kind, including translocations that relocate loci between chromosomes, are increasingly recognized as an important and underappreciated driver of animal genome evolution ^70^, making such an origin for IGH’s current chromosomal position plausible. If IGH’s present-day chromosome has carried this low-complexity character since that event, both consequences we describe (frequent inversion formation and the pseudogene pool that sustains gene conversion) would follow directly from that single historical event. We do not know the relative contribution of each, or whether both were in fact present from the outset; disentangling this will require further comparative and molecular work.

### V(D)J recombination mechanism predicts germline gene content and functional gene positioning at IGH and IGL

This difference in strand organization has direct implications for the V(D)J recombination mechanism employed at each locus. When a V gene and the (D)J segment share transcriptional orientation, rearrangement proceeds by deletion, excising the intervening sequence; when they are inverted relative to one another, it proceeds by inversion, which retains that sequence ^2,71^ (Figure 4). In birds, strand orientation therefore predicts the likely mechanism at each locus: at IGH, where V genes are distributed across both strands, both deletion and inversion are available, whereas at most IGL loci, where V genes lie predominantly on one strand, deletion is expected to predominate.

**Figure 4:**
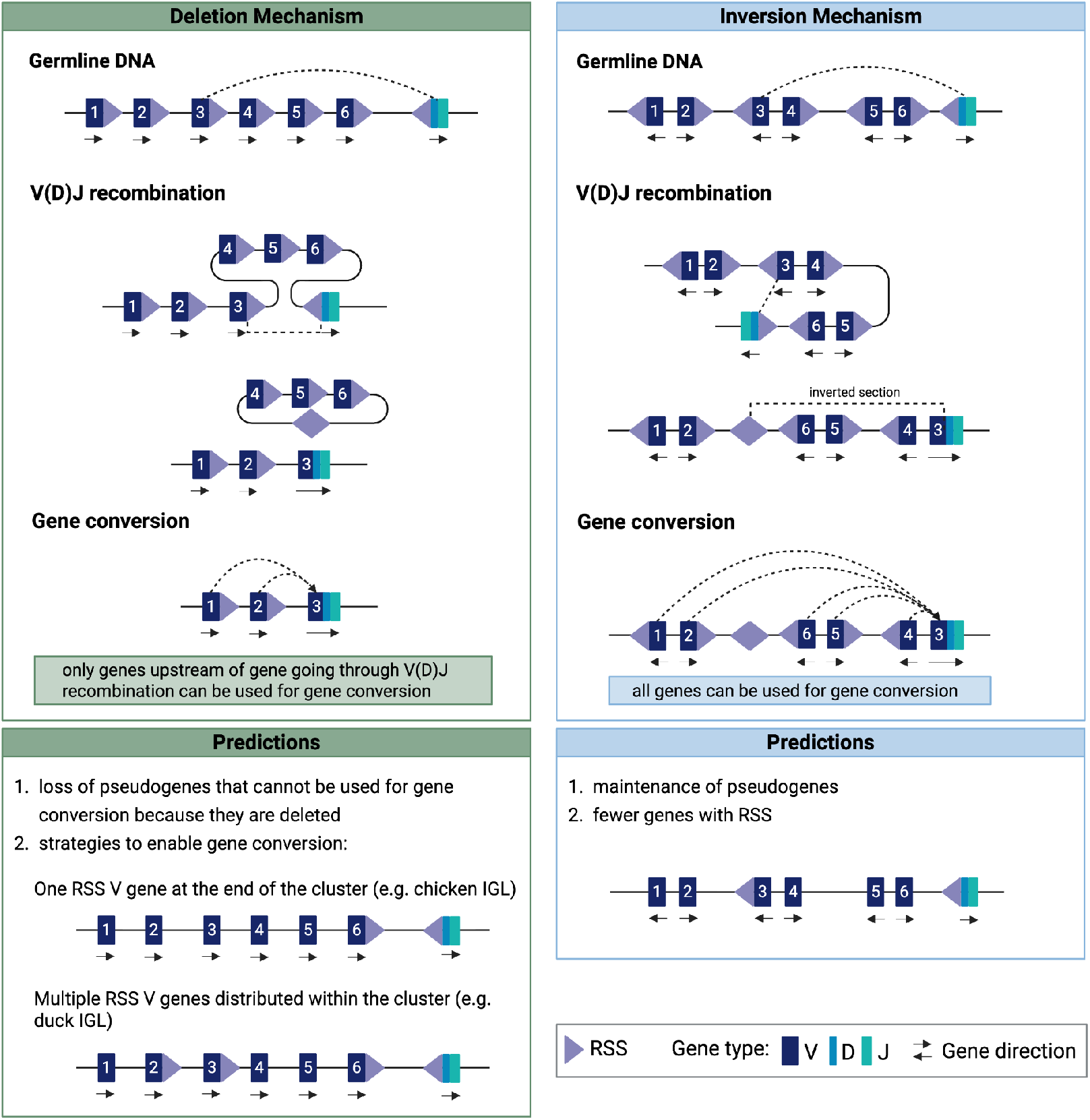
V(D)J recombination mechanisms and their impact on gene conversion Two different types of mechanisms are used in V(D)J recombination depending on the orientation of the genes and their recombination signal sequences (RSS): Deletion Mechanism (green): When the V gene and the (D)J segment share transcriptional orientation, their RSSs face each other. Double-strand breaks are made between each gene and its RSS, the V and (D)J segments are joined, and the intervening DNA is excised as a circle. In the example shown, V genes 4–6 lie between the rearranging gene and (D)J and are lost, leaving only genes 1 and 2 available as gene-conversion donors. Inversion Mechanism (blue): When the V gene and the (D)J segment are in opposite orientation, their RSSs point the same way. Breaks are again made at each RSS, but the intervening DNA is inverted rather than excised, so all germline genes are retained and remain available as donors after recombination. Note that this recombinatorial inversion is distinct from the germline inversions that characterize the avian IGH locus; the gene arrangement shown is schematic and illustrative, not a depiction of any specific locus.

Because the deletion mechanism excises every V gene between the rearranging gene and (D)J, any pseudogene downstream of the functional gene is lost and cannot act as a donor (Figure 4). Since only pseudogenes on the same chromosome contribute to gene conversion ^34,36,70^ a deletion-based locus can preserve its donor pool in only two ways. Either a single functional gene sits at the J-proximal terminus of the cluster, leaving all pseudogenes upstream and intact, or multiple functional genes are distributed across the locus so that donors remain available wherever rearrangement occurs. The former is the chicken IGL configuration, with its single terminal IGLV gene ^34,68,69^; the latter is seen in Pekin ducks, which carry multiple functional IGLV genes ^72^. The inversion mechanism imposes no such constraint: because the intervening sequence is retained, all pseudogenes remain available as donors regardless of position (Figure 4).

These mechanisms therefore predict contrasting locus architectures: IGL, rearranging by deletion, should carry few total genes but a higher proportion of functional RSS-associated genes, with any single functional gene biased toward the terminal position. IGH, where the inversion mechanism relaxes both constraints, should carry many pseudogenes, few RSS-associated genes, and no positional bias in the functional genes it does retain.

### Cross-species data confirm the predicted differences in gene content and functional-gene positioning

Before these predictions could be tested, RSSs had to be identified across the dataset to determine likely functional genes. Canonical mammalian RSS motifs failed to detect IGH RSS in essentially any bird, including close relatives of the chicken; only a dedicated discovery strategy built on the chicken motif with relaxed thresholds recovered candidates (see Methods, Supplementary Section 8). That RSS is conserved enough to be found even in non-avian reptiles ^3^ have diverged so far in birds is itself further evidence that the avian IG system has followed a distinct evolutionary trajectory.

Testing our previous predictions requires knowing which end of each V cluster is J-proximal. At IGL this is straightforward: because nearly all V genes share one strand, J lies downstream of that strand. At IGH, where genes occupy both strands, we instead used D gene annotations to orient the locus and locate the J-proximal terminus (see Methods; Supplementary Section 9). We then compared total and RSS-associated gene counts (see Methods, Figure S8.1, Figure S8.2) across IGH and IGL in all annotated species. Our findings are consistent with the predicted pattern: IGL loci carry significantly more RSS-associated genes and far fewer total genes, indicating a limited pseudogene pool, while IGH loci show the opposite trend, with few RSS-associated genes but large numbers of pseudogenes (Figure 5A).

**Figure 5:**
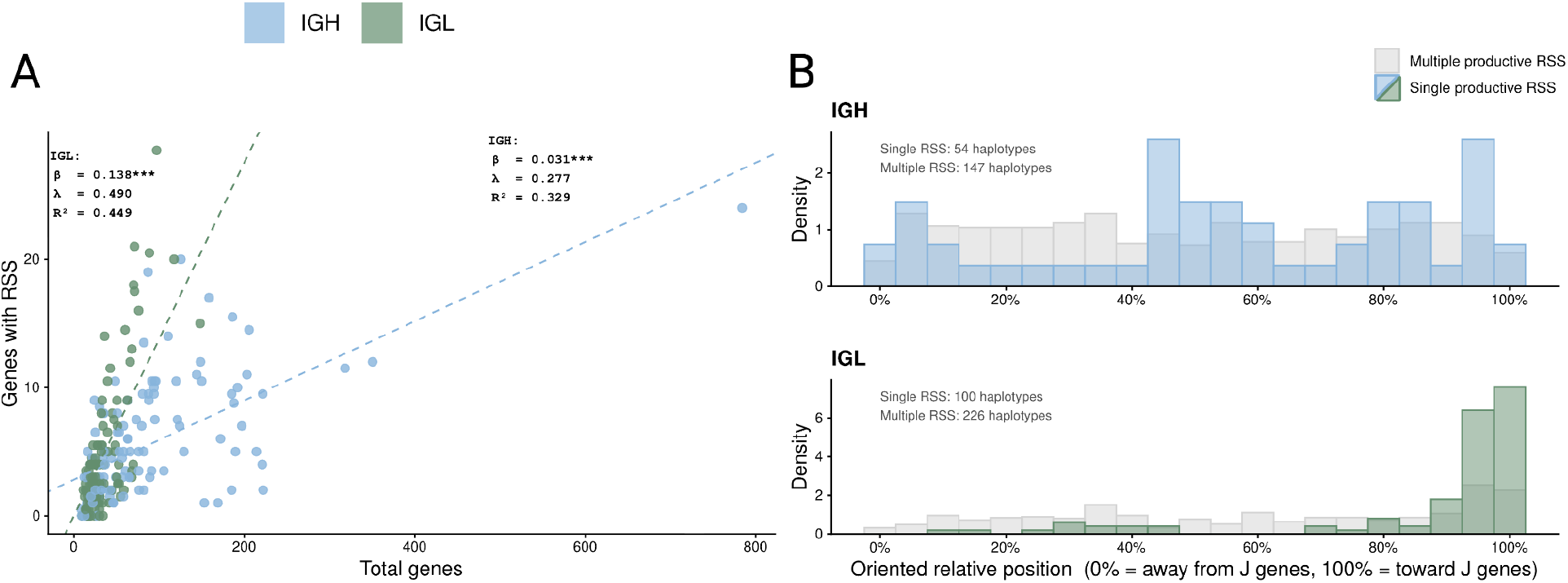
A) Number of total genes compared to the number of genes with an RSS per species. The correlations were calculated using *phylolm* ^82,83^ B) Relative position of RSS genes. We differentiate between haplotypes that have multiple productive genes with RSS (grey) and haplotypes with only one productive gene with RSS (IGH:blue, IGL: green)

We further analyzed the positions of RSS-associated genes within the V gene cluster, distinguishing between haplotypes with a single RSS-associated gene and those with multiple. For IGL haplotypes carrying only a single RSS-associated gene, the predicted positional bias is clearly supported: these genes show a significantly higher density at the terminal end of the V gene cluster (Figure 5B). Where multiple RSS-associated IGLV genes are present, a positional preference toward the terminal end of the cluster is still detectable but is considerably weaker (Figure 5B), consistent with the expectation that having multiple functional genes distributed across the locus relaxes the selective pressure for terminal positioning (Figure 5). In IGH, no strong positional bias is observed regardless of RSS gene count, though haplotypes with a single RSS-associated gene show an enrichment toward the middle of the cluster and around the 95th percentile position (Figure 5B). These results support our hypothesis that the deletion and inversion recombination mechanisms, in combination with gene conversion, have shaped the contrasting gene repertoire architectures of avian IGH and IGL. It is worth noting that although the inversion mechanism renders the position of the functional gene inconsequential for pseudogene retention, the strand distribution observed in avian IGH also permits the deletion mechanism to operate. This may explain why in the chicken the functional gene is found at the terminal position of the IGHV gene cluster, as this positioning would be advantageous under the deletion mechanism regardless of whether inversion-based recombination also occurs.

### Expressed repertoire data support gene conversion as the primary diversification mechanism

While germline analysis provides insight into locus architecture and gene content and allows us to generate hypotheses about how diversity is created, only a joint analysis of the germline and the expressed repertoire from the same individual can directly reveal gene conversion events and identify which genes are actually functional; i.e., the use of the same avoid the confounding effect of population variation at the germline alleles. To our knowledge, no such dataset exists for any bird aside from the chicken, which we already demonstrated appears to be an outlier among birds in features like the strand bias of IGL genes. To this end, we generated matched whole-genome and whole-blood Iso-seq (PacBio Kinnex) sequencing data from a wild-caught Red-winged Blackbird (*Agelaius phoeniceus*, Ithaca, NY). To confirm that this assembly named bAgePho2 provided a reliable basis for the analyses that follow, we assessed read alignment quality specifically at the IGH and IGL loci using CloseRead^13^, finding no evidence of the mapping quality, coverage, or mismatch anomalies that typically indicate misassembly at either locus (Figure S10.1). After filtering over 91 million long-read transcripts to a confident set of IG-derived sequences (see Methods), we recovered 678 transcripts aligning to both loci. On the primary haplotype, IGH reproduced the pattern established across our cross-species dataset: the locus is characterized by inversions (Figure S10.2) and has a large gene cluster (103 V genes) in which only a small minority carry an RSS (25 genes) (Figure 6A). At the IGL locus we found 22 V genes, only 2 of which carry an RSS (Figure 6B). On the alternate haplotype, only the IGL locus was recovered, with a matching gene count; IGH could not be confidently assembled on this haplotype (see Methods). Notably, no IGH J gene could be located in this assembly, despite an extensive search of the region downstream of the V array, indicating that transcript-to-gene assignment for IGH necessarily relies on direct alignment to V genes alone rather than full VDJ classification.

**Figure 6:**
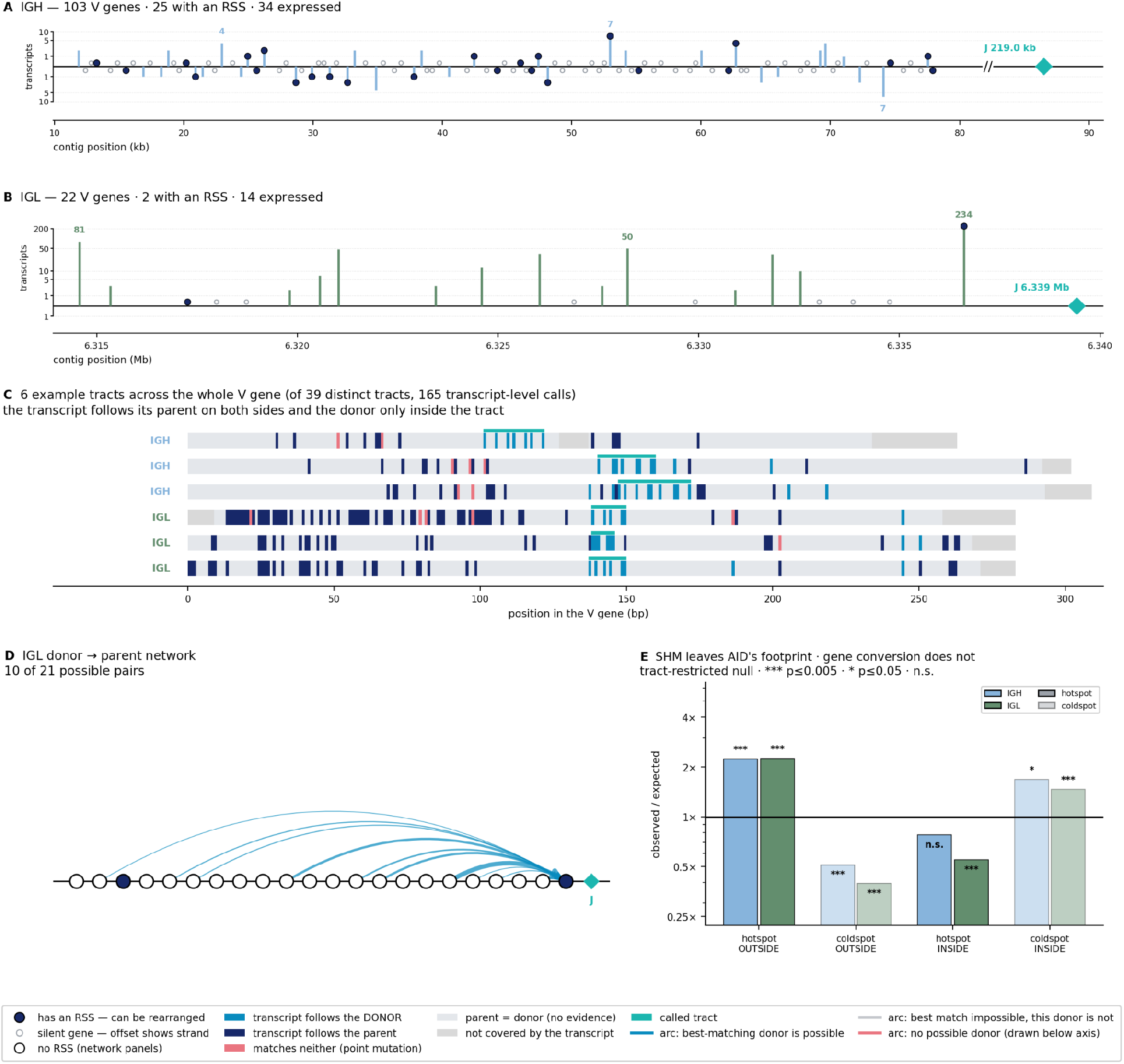
Gene conversion in the Red-winged Blackbird. A. IGH V gene annotations; stem height is transcript count (log), direction is strand, navy blue dots mark an RSS, open circles on the zero line mark genes that we found in the germline but were not expressed in the Iso-seq data. 25/162 IGH carry an RSS. 35/162 are expressed. Genes that are expressed but do not have an RSS are likely either donor genes that are very similar to a parent gene or they are parent genes with an unfamiliar RSS. Teal diamond, J B. IGL V gene annotations 2/23 IGL V genes carry an RSS. 15/23 are expressed. details as in A) C. Six example tracts across the whole V gene, three per locus, of 39 distinct tracts from 165 transcript-level calls: navy where the transcript follows its parent, teal the donor, rose neither, pale grey where parent and donor agree. Teal bar, called tract. The transcript follows its parent on both flanks and the donor only inside the tract. D. IGL donor → parent arcs, width is proportional to the number of supporting tracts; 10 of 22 possible pairs, all converging on the single functional gene. E. Differences outside tracts are hotspot-enriched and coldspot-depleted in both loci (all p = 0.002), the hypermutation signature; inside tracts the pattern is absent or reversed (IGL ×0.55 and ×1.47, both p = 0.002; IGH ×0.78, n.s., and ×1.68, p = 0.014), as predicted if those differences were copied from a donor and therefore lie wherever the donor happened to differ from the parent, independently of where AID acted. Enrichments are relative to 1,000 tract-restricted permutations, with inside-tract differences counted once per clone per tract. *** p ≤ 0.005; * p ≤ 0.05; n.s., not significant.

Restricting to genes with robust transcript support (see Methods), only 19 of the RSS-associated IGH genes and a single RSS-associated IGL gene were confirmed as expressed. The expressed IGL gene occupies the position closest to J within the V cluster, consistent with our deletion mechanism model, which predicts that the sole gene able to survive rearrangement should occupy this terminal position. In IGH, where the inversion mechanism should render position largely inconsequential for pseudogene retention, expressed genes show no comparable positional bias. The germline architecture and the expressed repertoire thus converge on the same conclusion from independent lines of evidence.

To test for gene conversion, we searched for stretches of transcript sequence that matched a donor V gene more closely than the expressed parent gene at multiple donor-diagnostic positions, requiring supporting positions to fall within 5 bp of one another to qualify as a contiguous tract rather than scattered chance agreement (Figure 6C; see Methods). Calibrating a conservative detection threshold against two independent controls, the rate of spurious tracts expected under permutation, and whether candidate tracts inadvertently capture ordinary mutational hotspots (detailed below), this approach identified 122 candidate conversion tracts in IGL (23.1%) and 28 in IGH (20.6%). Interestingly, the majority of tracts are located in the complementarity-determining regions (CDRs) both in IGH and in IGL (Supplementary Section 11, Figure S11.1) as has been found in a previous study in chicken^69^. There is also a peak in the framework region 3 (FR3) in IGL.

The IGL locus provides a particularly stringent test of the model, as its architecture eliminates a key potential confound: with a single functional gene positioned at the deletion-safe terminus of the cluster, no candidate donor gene is ever excised by recombination, and donor availability is therefore never in question. Every one of the IGL tracts we identified has a topologically valid donor, and tracing each tract to its most probable source reveals ten distinct donor genes, all converging on the same single expressed parent (Figure 6D). This is the expected signature of gene conversion as characterized in the chicken, now demonstrated directly in a second avian species.

The IGH locus presents a more complex picture, and the source of that complexity is itself informative. Because IGH genes rearrange by either deletion or inversion depending on strand orientation, a subset of candidate donor assignments implicate genes that should, in principle, have already been excised at the time of the corresponding rearrangement, an apparent violation of the model (see Supplementary Section 11, Figure S11.2). We interpret this not as a misspecification of the underlying mechanism but as a limitation of donor attribution under conditions of extreme sequence similarity: with 103 highly homologous candidate donors, the true source of a given tract is frequently statistically indistinguishable from one or more donors that are topologically implausible. The model can detect that a conversion event has occurred with considerably more confidence than it can identify which of several near-identical genes supplied it, a limitation that follows directly from the same large, homogeneous pseudogene pool that makes gene conversion effective at this locus in the first place.

The most decisive evidence does not depend on donor identity at all. Somatic hypermutation and gene conversion are both initiated by the enzyme AID, but the two processes are expected to leave distinguishable molecular signatures: hypermutation should cluster at AID’s preferred target motifs ^73–77^, whereas gene conversion should introduce whatever sequence differences happen to exist between donor and parent, independent of AID’s targeting preference. Comparing sequence differences outside versus inside called conversion tracts confirms this prediction directly. Outside tracts, differences are significantly enriched at AID hotspot motifs (WRCY/RGYW) and depleted at coldspots (SYC/GRS) in both loci (IGH: 2.23× and 0.51×; IGL: 2.24× and 0.40×; all p = 0.002), consistent with active somatic hypermutation. Inside tracts, this signature is absent or reversed: in IGL, tract-internal differences were depleted at hotspots (0.55×) and enriched at coldspots (1.47×; both p = 0.002) the pattern expected if these differences were copied from a donor sequence rather than introduced by AID at its preferred sites. In IGH, the same reversal was present at coldspots (1.68×, p = 0.014), while the hotspot shift was smaller and did not reach significance (0.78×, n.s.) (Figure 6E). Together, these results provide direct molecular evidence that gene conversion, and not solely somatic hypermutation, actively diversifies the expressed IG repertoire in a bird other than the chicken. Scoring transcripts against different germline references introduces false parent-gene assignments and downstream errors in tract detection and donor attribution (see Supplementary Section 12), underscoring the importance of matched individual-level genome and transcriptome sequencing for this kind of analysis.

## Discussion

The cross-species comparison of avian IG loci presented here reveals both a surprising degree of conservation and a remarkable degree of diversity, and underlines how little we have known about avian IGs outside of a single model organism. Almost all birds we analyzed share similarly short locus lengths, a lack of strand bias in IGH, and the newly described inversion architecture of the IGH locus, yet there are substantial differences in inversion burden and gene counts across species. Our population-level analysis adds another layer to this picture: even within a species, IGH locus architecture can vary considerably between individuals, with the contrast between the small, isolated Island Scrub Jay population and the widely distributed Woodhouse’s Scrub Jay illustrating how demographic history leaves a detectable imprint on IG locus structure. Together, these observations underscore how dynamic the IG loci are, and serve as a reminder that characterizing IG diversity from a single individual or a handful of haplotypes per species will inevitably underrepresent the true extent of variation.

Our confirmation that all 122 annotated bird species lack an IGK locus establishes its loss as a universal avian feature. This, alongside the known reduction to a single functional IGHV gene in chickens and the simultaneous rather than sequential rearrangement of IGH and IGL ^27,69,78,79^, may reflect a broader simplification of the V(D)J rearrangement step in birds. In mammals, combinatorial diversity across many VH genes and two light chain loci makes the rearrangement step itself a major source of antibody diversity^80^, and the ordered developmental checkpoint, in which heavy chain rearrangement precedes the kappa light chain and then lambda light chain as a fallback, serves to quality-control and amplify successful rearrangements before committing to a light chain ^71^. In birds, however, we show that diversity is generated almost entirely by gene conversion rather than at the rearrangement step, extending a process known to occur in the bursa of Fabricius in the chicken more broadly across birds. This may have removed the selective pressure to maintain a large VH repertoire, two light chain loci, and an ordered rearrangement checkpoint independently. Which of these features was lost first, and whether they are mechanistically linked or independent consequences of the shift to gene-conversion-based diversification, remains an open question.

The connection between inversions and gene conversion we propose here offers a coherent explanation for an otherwise puzzling genomic feature. Repeated inversions can cause instability of chromosomes ^59,64–66^ but at avian IGH, they appear to be maintained by selection because they continuously regenerate the raw material required for gene conversion: copies of existing genes on the opposite strand from the functional gene, with sufficient sequence similarity to serve as conversion donors. The strong positive correlation between inversion count and V gene number, and the observation that the majority of IGH genes in a given species reside on inverted segments, suggest that inversions are a primary mechanism for expanding the pseudogene pool rather than an incidental byproduct of some other process. The relative contribution of the mutation-rate and tolerance effects proposed above (see Results), and how they interact, will require further molecular work to resolve.

The restriction of the inversion pattern to IGH in most species, despite gene conversion operating at both IGH and IGL in the chicken, is explained above by chromosomal context (see Results). The independent acquisition of IGL inversions in landfowl and in the clade comprising hummingbirds, nightjars and swifts is particularly intriguing under this model, as it implies that something specific to these lineages tips the cost-benefit balance in favor of tolerating inversions at IGL as well. Whether this reflects differences in pathogen exposure, or some other lineage-specific factor, remains an open and compelling question.

The chicken has long served as the default model for avian immunology, but until now it was unknown which of its unusual IG features were general avian adaptations and which were peculiarities of a single domesticated species. Our results resolve this. Using matched germline and transcriptome sequencing from a wild Red-winged Blackbird, we provide the first direct molecular evidence that gene conversion diversifies the expressed antibody repertoire in a bird other than the chicken, and, extending this across 122 species, show that the underlying strategy, a restricted set of functional V genes sustained by a continuously regenerating pseudogene pool, is general across birds rather than chicken-specific. The chicken is nonetheless not fully representative: its lack of directional preference in IGL gene orientation, driven by inversions, is shared with landfowl but otherwise unusual among birds. Gene conversion itself, in other words, appears to be an ancestral, shared avian strategy, while the inversion-driven architecture that sustains it at IGH is a more variable, independently evolving trait, underscoring that any single species, however well studied, risks missing the true breadth of evolutionary solutions to generating immune diversity.

## Methods

### V Gene Annotation

The large-scale annotation of IG loci across VGP species required the development of new methods and databases capable of handling the breadth of species represented. Specifically, the IgDetective tool^5^ was expanded to enable the annotation of avian V genes. IG loci and genes were annotated using the avian database of IgDetective, following the approach described in Formenti et al. 2026^3^. We opted to use the database from VGP iteration 2, as subsequent iterations showed a marked increase in the number of contigs containing annotated genes while failing to recover the original reference genes from the chicken, suggesting a loss of annotation specificity. This conservative choice produced the most consistent and biologically sensible results across species. We also made sure to only include birds for which we have both IGH and IGL annotations.

### Dotplots

We generated the dotplots shown in Figure 1D, Figure 2 and supplemental figures using Patchworkplot ^84^. It visualizes the alignment of two sequences (in our case of a locus to itself): Each axis represents one of the two sequences being compared, with dots placed at positions of sufficient similarity, such that diagonal lines indicate conserved sequence and inverted diagonals indicate inversions. We used the parameters --show-annot --lower --lwidth 2 --cmap viridis --min-pi 75 --transparent. In Figure 1D and Figure S4 we used --min-len 1500 and in Figure 2 --min-len 10000 to show a broader pattern. For all other supplemental figures we used --min-len 15000. Note that the relative sizes of the dotplots in Figure 1D are not proportional to the actual sizes of the respective loci, which vary across species.

### V Gene Filtering

The IgDetective annotation pipeline occasionally produces false positive annotations of sequences unlikely to represent true IG genes. To address this, we applied two additional filtering steps to all annotated genes: The first filter operates at the contig level. In several cases, multiple contigs containing potential IGH or IGL genes were identified within a single haplotype. Where one contig harbored substantially more genes than the others, the genes on the smaller contigs were considered likely false positives. However, in cases where multiple contigs had comparable gene counts, excluding all but the largest would risk discarding potentially functional genes. We therefore filtered out all genes on contigs whose gene count was less than 20% of that of the highest-gene-count contig within the same haplotype, preserving genes from all larger contigs while removing only those on the smallest ones. The second filter operates at the gene level. A proportion of annotated sequences were either too short or too low in complexity to plausibly represent functional IG genes. We therefore removed all sequences shorter than 250 bp, as well as all sequences in which 70% or more of the content consists of homopolymers.

### RSS discovery

Since the RSS related to the functional chicken IG genes differ from the canonical mammalian motifs, IgDetective, whose search procedure is based on mammalian motifs, does not detect these RSS. To identify RSS in birds more broadly, we developed a dedicated RSS discovery procedure to supplement IgDetective. This procedure takes all annotated IG genes as input and searches for candidate RSS sequences using a database of bird heptamer and nonamer motifs. However, given the limited scope of prior studies, this database contains only two known bird RSSs within IGHV: the chicken RSS and the RSS associated with the most highly expressed gene in a tufted duck Iso-seq dataset ^85^. This issue of limited scope is present within IGLV as well, but the RSS related to the functional gene within this locus differs much less from the mammalian motif than in IGHV. Due to this, we opted to use a dataset of only one RSS for this locus, with that being the most commonly seen motif within IGLV in humans. Because direct motif matching, as used by IgDetective, would therefore fail to capture the full diversity of avian RSS, we instead searched for indirect matches, identifying candidate sequences that resemble known motifs within a defined number of base pair differences. Through comprehensive testing of various thresholds, we settled on allowing up to 2 bp differences for heptamers and up to 6 bp differences for nonamers. Using these thresholds, we searched 0 to 2 bp downstream of every annotated gene for a candidate heptamer and nonamer, flagging any gene where both were present as a potential RSS-associated gene. Within these candidate genes, we first applied the filtering procedure described above, and we then analyzed the remaining potential heptamers and nonamers; quantifying these sequences independently of each other through the number of appearances in different haplotypes. Using these quantifications, we removed heptamers that appeared less than 10 times and nonamers that appeared less than 5 times across the dataset, and used the remaining top candidates to identify bird heptamer and nonamer motifs within IGHV and IGLV (Figure S8.1 and S8.2).

### Sample collection and nucleic acid extraction

A single male Red-winged Blackbird (Agelaius phoeniceus) was captured by mist-netting on a defended territory in Ithaca, NY; sex was determined in the field from plumage (glossy black with the characteristic red-and-yellow epaulets of adult males, in contrast to the streaked brown plumage of females) and was consistent with the bird’s aggressive, territorial response to conspecific playback. Sex was further confirmed genomically by two independent tests: aligning both bAgePho2 haplotypes to the assembly of a conspecific female (bAgePho1.hap1) recovered a full-length Z chromosome in both haplotypes, with the W chromosome absent except for the pseudoautosomal region; and mapping a 5% subsample of the bird’s own HiFi reads to the primary assembly showed the Z chromosome at approximately 0.99× autosomal depth, indicating a diploid (ZZ) rather than hemizygous (ZW) genotype. A blood sample was collected from the brachial vein and preserved in DNA/RNA Shield (Zymo Research). High molecular weight genomic DNA was extracted using the Quick-DNA HMW MagBead Kit (Zymo Research), quantified using a Qubit fluorometer (Thermo Fisher Scientific), and assessed for integrity using the Femto Pulse System (Agilent). Whole-blood RNA was extracted using the Quick-DNA/RNA Miniprep Plus Kit (Zymo Research) following the manufacturer’s protocol for whole blood, quantified using a SpectraMax QuickDrop (Molecular Devices), and assessed for integrity using a Fragment Analyzer (Agilent). Library preparation and sequencing were performed at the Genomics Resources Core Facility, Weill Cornell Medicine. The whole-genome sequencing library and the Iso-seq library were each sequenced on a separate PacBio HiFi SMRT Cell.

### Red-winged Blackbird whole genome assembly

We assembled the genome from PacBio HiFI reads only using hifiasm ^86^ (v0.25.0-r726) in primary-alternate mode. The input comprised 8,203,858 HiFi reads totalling 160.6Gbp (mean read length 19.6kbp) giving approximately 128x coverage of the assembled genome. Default error correction (three rounds) and purge-duplication settings were used. The primary assembly is 1.25Gbp in 264 contigs, with N50 of 65.1Mbp (L50=7) and a longest contig of 155.1Mbp. The associated alternate assembly is 1.19Gbp in 698 contigs, with N50 of 26,9 Mbp

### Iso-seq library generation and initial processing

We generated PacBio Kinnex (MAS-8) full-length cDNA sequencing data from whole blood RNA of the same Red-winged Blackbird individual (bAgePho2) used for the whole-genome assembly, enabling matched germline and transcriptome analysis. Kinnex reads, which concatenate approximately eight cDNA molecules per sequencing read, were split into individual segments using `skera split` against the MAS-8 adapter set. Reads were then trimmed of 5’ and 3’ cDNA primers and oriented using `lima --isoseq`, and full-length non-concatemer (FLNC) reads were generated using `isoseq refine --require-polya`, retaining only reads with a complete cDNA structure from 5’ cap to polyA tail. This yielded 91,291,873 FLNC reads.

Because this is a whole-transcriptome library from blood and only IG-derived transcripts were of interest, FLNC reads were prescreened by alignment to the annotated V gene set prior to clustering, retaining reads with ≥70% identity over ≥150 bp (4,767 reads). These were clustered into consensus transcripts using `isoseq cluster2`, yielding 709 transcripts, of which 676 were IG-derived (109 IGH, 567 IGL) based on the prescreen. Transcript-level IG status was confirmed using two independent approaches: direct alignment to the V gene set via minimap2, and VJ classification using immunotools’ `vj_finder`. The two methods agreed on 509 transcripts; minimap2 uniquely identified an additional 167, and immunotools an additional 2, for a combined total of 678 IG transcripts. Because no IGH J gene could be identified in the bAgePho2 assembly (see below), immunotools, which requires both a V and J match, could not classify IGH transcripts, and IGH transcript identification therefore relied on the minimap2 alignment approach alone.

Transcripts were aligned to the V gene reference set of the primary haplotype using minimap2 (`-x splice:hq --cs`), and the resulting `cs` tag was used to derive base-level differences between each transcript and its best-matching germline V gene. All identity and difference calculations excluded the terminal 20 bp of each V gene, since this region is truncated and modified by non-templated nucleotide addition during V(D)J junction formation independent of any post-recombination diversification process.

### Gene and transcript expression classification

A V gene was classified as RSS-associated based on the curated RSS annotation described above (see RSS discovery). A gene was classified as expression-confirmed when supported by ≥2 transcripts matching it at ≥98% identity over ≥200 bp, excluding the VDJ junction region. A small number of genes showed confirmed expression despite lacking an annotated RSS. We flagged these separately, since there are two likely explanations: either the gene does carry a functional RSS that our discovery procedure failed to detect because it diverges too much from the known motifs, or the gene is not itself rearranged but is a donor whose sequence so closely resembles the true rearranged parent gene that transcripts derived from the parent were mistakenly matched to the donor instead.

### Gene conversion tract detection

For each expression-confirmed parent gene, we identified transcript positions differing from the parent germline sequence and asked whether each differing base matched the corresponding position in an alternative germline V gene (a candidate donor), defining such positions as donor-diagnostic. Contiguous stretches of donor-diagnostic positions, defined as positions falling within 5 bp of one another, were called candidate gene conversion tracts. This contiguity requirement was necessary to distinguish genuine templated tracts from spurious agreement scattered across a transcript by chance; without it, apparent “tracts” had a median span of 53 bp supported by a median of only 7 positions.

The minimum number of supporting donor-diagnostic positions (m) required to call a tract was calibrated independently for each locus using two controls. First, a permutation null was constructed by redistributing each transcript’s observed differences to random positions while preserving their number and identity, over 20 replicates; this establishes the false-tract rate expected under no true tract structure. Second, the AID mutational spectrum test (below) was used to detect thresholds too permissive to exclude clustered somatic hypermutation, since an overly loose threshold causes tract-internal differences to carry an AID-hotspot signature they should not have if the tracts are genuine templated events. Based on these controls, we set m ≥ 5 for IGL and m ≥ 6 for IGH; both thresholds fall comfortably above the level at which the permutation null approaches zero false tracts, and the AID-spectrum contrast (below) was confirmed to be stable across m = 5–8 in both loci, indicating that the specific threshold chosen does not drive the result.

Where multiple candidate donor genes shared identical diagnostic positions for the same tract, all were retained as competing explanations of a single conversion event, and the candidate with the greatest number of supporting positions was designated the primary donor.

### Cis-acting donor availability and locus topology

Gene conversion was modeled as strictly cis-acting, requiring the donor to reside on the same chromosome as the recombining locus. Whether a given candidate donor remained physically available at the time of V(D)J recombination was determined by the inferred recombination mechanism for that parent gene, based on the relative strand orientation of the V gene and the J segment (see Results, VDJ recombination mechanism). Under the deletion mechanism, all V genes located between the rearranging V gene and J are excised and lost; under the inversion mechanism, the intervening sequence is inverted but retained, and all donors remain available. For each tract, we evaluated whether its assigned donor(s) were topologically consistent with the recombination mechanism of the corresponding parent gene; tracts for which every candidate donor had necessarily been excised prior to rearrangement were flagged as topologically impossible. Because IGL in this species rearranges exclusively via the deletion mechanism from a single, J-proximal functional gene, no IGL donor is ever excised, and this control has no discriminating power in that locus; it is informative primarily for IGH, where both mechanisms operate.

### AID hotspot spectrum analysis

To distinguish somatic hypermutation from gene conversion at the level of individual transcript-to-germline differences, we classified each difference position according to whether the corresponding germline parent sequence matched a canonical AID hotspot motif (WRCY or its reverse complement RGYW) or coldspot motif (SYC or its reverse complement GRS), using IUPAC ambiguity codes (W = A/T, S = G/C, R = A/G, Y = C/T). The motif was always evaluated on the parent germline sequence, not the transcript, since the relevant question is whether AID would have targeted that genomic position. For each locus and tract status (inside versus outside a called conversion tract), we computed the fraction of differences falling at hotspot and coldspot positions and compared this to a null distribution generated by 1,000 permutations in which each transcript’s observed mutation count was redistributed at random among the positions it covered. Two permutation frames were used: a gene-wide null, drawing from all positions in the gene, and a tract-restricted (strict) null, drawing separately from within-tract and outside-tract positions for each transcript. The strict null is reported as the primary result, since tract windows in IGH are compositionally unusual relative to gene-wide background (1.5-fold enriched for hotspot motifs, 2.4-fold depleted for coldspot motifs), and using gene-wide composition as a baseline for tract-internal differences would conflate composition with process. Two-sided empirical p-values were computed with a standard +1 correction, giving a minimum achievable p-value of 0.002 at 1,000 permutations. Enrichment was calculated as the observed fraction divided by the mean of the null distribution.

## Resource availability

### Lead Contact

Matt Pennell is the lead contact.

### Materials Availability

The new assembly (bAgePho2), the reads and the Iso-seq data for the Redwinged Blackbird can be found under BioSample SAMN62949155, BioProjects PRJNA1524053 (WGS reads and Iso-seq reads), PRJNA1524069 (primary assembly) and PRJNA1524068 (alternate assembly).

### Data and Code Availability

All the assembly data is publicly available through GenomeArk^87^. We used the species tree provided by the Phase I of the Vertebrate Genome Project ^3,88^.

The code for the annotation pipeline can be found at:

https://github.com/applied-phylo-lab/VDJ_Annotation_Pipeline. The Iso-seq pipeline can be found at:

https://github.com/applied-phylo-lab/isoseq_pipeline. All other code used for the analysis can be found at: https://github.com/applied-phylo-lab/Bird_IG

## Acknowledgments

We thank Corey Watson, William Lees, and Erik Enbody for insightful and spirited discussions on this work.

## Author contributions

Conceptualization: K.V., M.Pen., Y.S., Y.Z.

Methodology: K.V., D.H., A.Z., M.Pos., Y.S.

Data curation: K.V.

New Data Collection: M.R.C., L.C.

Data analysis and visualization: K.V., D.H., A.Z.

Writing – Original Draft: K.V., D.H., M.Pen.

Writing – Review & Editing: all authors

Funding Acquisition: M.Pen.

Supervision: M.Pen., Y.S., L.C., A.B.

## Declaration of generative AI and AI-assisted technologies

During the preparation of this work, the authors used Claude in order to refine the structure and phrasing of the text. After using this tool or service, the authors reviewed and edited the content as needed and take full responsibility for the content of the publication.

## Supplement

### Supplementary Section 1: Contig length across vertebrates

**Figure S1:**
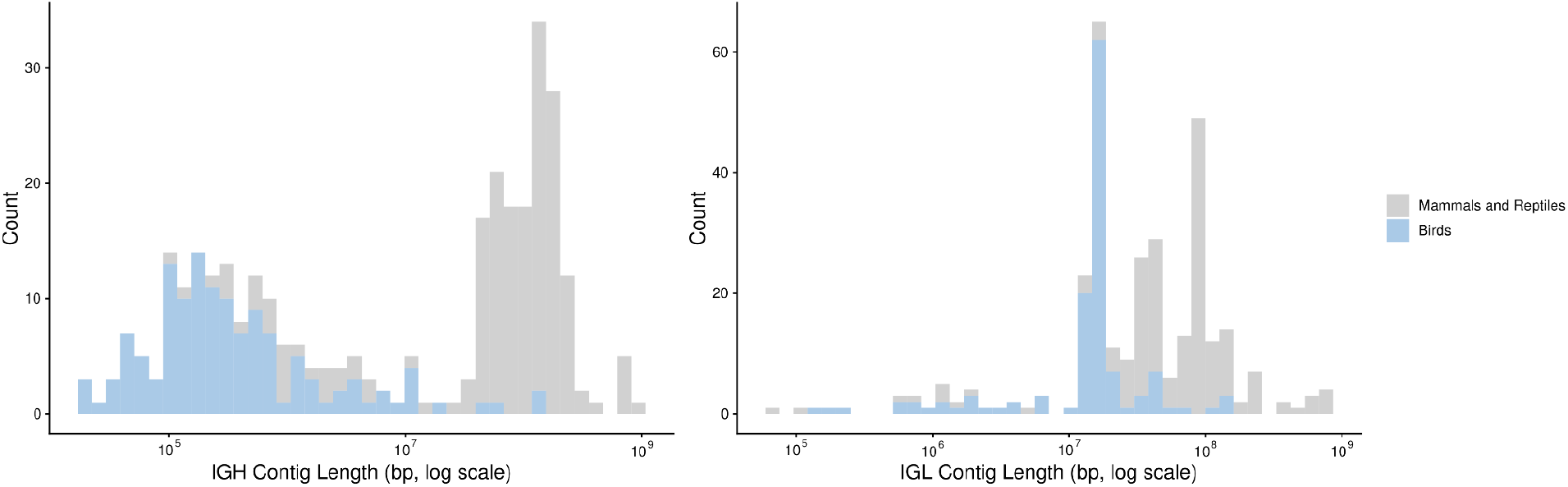
Contig length at the IGH and IGL loci in birds compared to other vertebrates Length (bp, log scale) of the contigs on which **(A)** IGH and **(B)** IGL are annotated, compared between birds (blue) and mammals and reptiles (grey) from the VGP dataset.

### Supplementary Section 2: Homopolymer enrichment and HiFi coverage dropout at IGH

#### Results

Beyond the breakage risk posed by inversions themselves, we asked whether the sequence context surrounding IGH independently contributes to assembly difficulty. The species for which we were able to identify sufficiently contiguous IGH-spanning scaffolds were predominantly assembled using combinations of PacBio HiFi with Hi-C or Oxford Nanopore Technology data. Scaffolds constructed using Hi-C scaffolding also contain many gaps filled with stretches of Ns, implying that HiFi sequencing of these regions is prone to coverage dropouts. This pattern points to intrinsic sequence properties of the IGH locus and its chromosomal context as a source of sequencing depth dropouts.

In most bird species, IGH resides on dot chromosomes — the smallest and most GC-rich members of the avian karyotype ^89–92^. Recent T2T assemblies of the zebra finch genome have shown that dot chromosomes are disproportionately enriched in minisatellite repeats and non-canonical (non-B) DNA motifs within their euchromatic compartments, and that these features are strongly associated with reduced PacBio HiFi sequencing depth: Formenti et al. reported a Spearman correlation of ρ = −0.69 between HiFi depth and minisatellite density across zebra finch dot chromosomes, with non-B DNA motifs being 56% more abundant in previously unassembled regions than in assembled ones ^3,93^. Whether the same properties underlie IGH assembly difficulty more broadly across birds had not been systematically examined.

To address this, we identified genome assemblies from 24 bird species spanning diverse avian orders for which a single haplotype provided a sufficiently contiguous scaffold to span the complete IGH or IGL locus together with at least 2 Mb of flanking chromosomal context. For each scaffold, we computed HiFi sequencing depth and a panel of sequence features — homopolymer richness, GC content, and the density of seven non-B DNA motif classes — in non-overlapping 4 kb windows, and assessed their association with coverage dropout using Spearman correlation meta-analysis across species (see Methods, Table S2.1).

Examination of individual assemblies revealed a consistent and visually striking pattern at IGH. In the representative example of the Greater scaup (*Aythya marila*; Figure S2A), the IGH-containing dot chromosome shows a pronounced reduction in HiFi coverage coinciding precisely with a sharp elevation in homopolymer richness. IGH locus is contained within this region of low HiFi coverage. No comparable signal is visible elsewhere on the same chromosome. The IGL-containing microchromosome, by contrast, shows uniformly high coverage throughout, with no elevation in homopolymer richness, a stark contrast that suggests the two loci occupy fundamentally different sequence environments despite both residing on microchromosomes.

These observations were confirmed quantitatively across all 24 species. Contigs containing IGH loci showed reduced HiFi sequencing coverage both within the locus and in its immediate flanking sequence. Within the IGH locus, normalised HiFi coverage was reduced to approximately 0.5 relative to the assembly mean haploid coverage; however, the most severe dropout with bins approaching near-zero normalised coverage was concentrated in the immediate flanking regions surrounding the locus boundary, with coverage recovering gradually with increasing distance (Figure S2B, left). This spatial pattern is consistent with the observation that the majority of avian IGH-containing contigs terminate at or near the locus boundary. No comparable dropout was observed at IGL loci, where coverage remained close to 1.0 throughout the locus and its flanking context (Figure S2B, right).

The homopolymer richness was elevated specifically in the IGH chromosomal neighbourhood and showed association with coverage dropout. Bins immediately flanking the IGH locus exhibited the highest homopolymer richness values in the dataset, consistently exceeding the background level of ∼0.25 typical of distal sequence, while in-locus bins showed intermediate elevation. At IGL, neither in-locus nor flanking bins showed comparable enrichment in homopolymer richness (Figure S2C).

Together, these results identify homopolymer richness as the primary sequence correlate of HiFi coverage dropout at avian IGH loci, with a stronger association than the non-B DNA motifs previously implicated in dot chromosome assembly difficulty ^3,93^. Notably, the most severe dropout is concentrated in the immediate flanking sequence rather than within the locus itself, which may explain why IGH loci are frequently recovered in assembled contigs even when the surrounding chromosomal sequence is not. IGL loci show no comparable dropout signal, consistent with their residence on longer microchromosomes with a distinct sequence composition.

**Figure S2:**
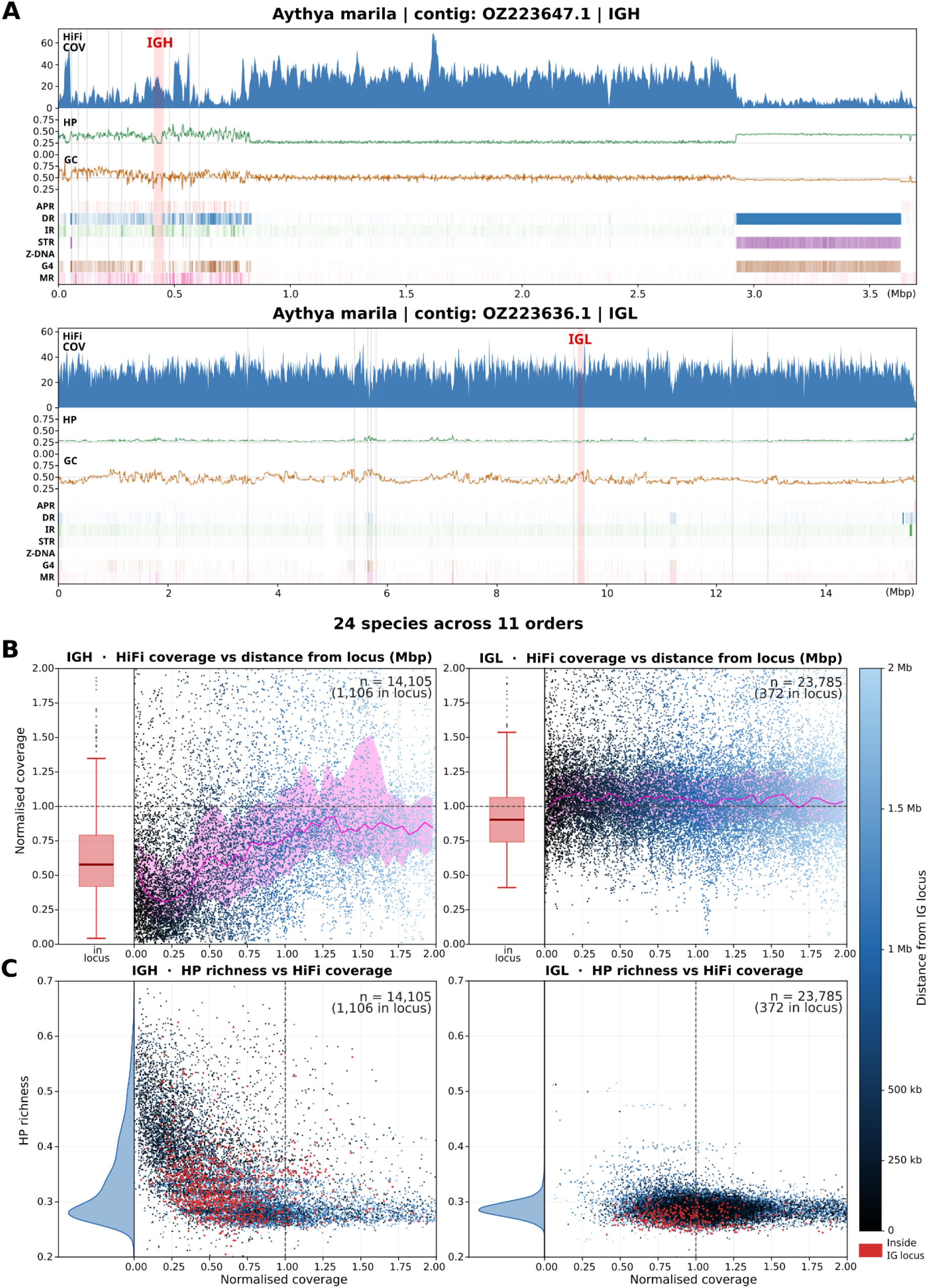
HiFi sequencing depth in IG loci neighborhood A) Genomic track plots for dot chromosome 28 (OZ223647.1) and microchromosome 17 (OZ223636.1) of Greater scaup (*Aythya marila*; genome assembly bAytMar2.hap1.1), containing the IGH and IGL loci, respectively, shown as a representative example. Tracks from top to bottom: HiFi sequencing depth (**HiFi COV**); homopolymer richness (**HP)**; fraction of bases in each bin covered by homopolymeric runs); GC content (**GC**); and density of non-canonical (non-B) DNA motifs: A-phased repeats (**APR**), direct repeats (**DR**), inverted repeats (**IR**), short tandem repeats (**STR**), Z-DNA (**Z-DNA**), G-quadruplexes (**G4**), and mirror repeats (**MR**). IG locus boundaries are highlighted by pink vertical shading; assembly gaps are indicated by gray vertical lines. B) Analysis of HiFi sequencing depth for the 2Mb flanking context of IGH (left) and IGL (right) loci across 24 avian species. For each locus, the left panel shows the distribution of in-locus HiFi sequencing depth normalised by the median haploid coverage for each species; the right panel shows normalised sequencing depth against distance from the locus boundary, with the magenta line and shaded band representing the median and interquartile range across species. All values are computed using 4 kb bins. C) Relationship between homopolymer richness and normalised HiFi sequencing depth for the same 4 kb bins as in (b), for IGH (left) and IGL (right) loci. Points are colored by distance from the locus boundary; bins overlapping the locus are highlighted in red. Marginal density plots show the distribution of homopolymer richness across all bins.

#### Methods

##### Species selection for assembly evaluation

We identified avian HiFi datasets by searching NCBI SRA and the GenomeArk repository ^87^ for PacBio HiFi (CCS) reads matched to published diploid genome assemblies. From an initial survey of 122 bird assemblies 24 species had publicly available HiFi read data, annotated immunoglobulin loci, and at least one contig spanning the complete IG locus with ≥2 Mb of assembled flanking sequence. Table S2.2 lists representative contig identifiers and lengths. For each species and each IG locus (IGH, IGL), we retained one representative contig. Where the diploid assembly carried two haplotypes on distinct contigs, we selected the contig annotated as the primary haplome unless the alternate contig was substantially more contiguous.

##### HiFi read alignment and coverage computation

We aligned HiFi reads to the corresponding diploid genome assembly using minimap2 ^94^ (v2.24-r1122) with the -ax map-hifi preset. We coordinate-sorted and indexed filtered alignments with samtools sort and samtools index (SAMtools v1.13) ^95^, retaining only primary alignments and discarding supplementary and secondary alignments. We estimated median sequencing depth for each assembly using samtools depth; depths ranged from 18× to 122× across species. We computed coverage across IG locus contigs in non-overlapping 4 kb bins and normalised per-bin coverage to the species-wide median, so that a value of 1.0 corresponds to the genome average for that species.

##### Sequence feature computation

We computed GC content per 4 kb window as the fraction of G and C bases among all non-N bases, and homopolymer richness (HP) per 4 kb bin as the fraction of bases identical to their immediate predecessor in the sequence, such that values near zero indicate alternating sequence and values near 1 indicate near-uniform mononucleotide composition.

We annotated non-B DNA structural motifs using the non-B GFA tool (v2) ^96^ across seven classes: A-phased repeats (APR), direct repeats (DR), inverted repeats (IR), short tandem repeats (STR), Z-DNA-prone sequences, G-quadruplex motifs (G4), and mirror repeats (MR). We applied the tool to FASTA sequences of each representative contig and computed per-bin density for each motif class as the number of annotated motif bases per 4 kb.

##### Statistical analysis for sequence features

We pooled all 4 kb bins from the 24-species dataset into a single dataset per locus (IGH, IGL) and window type (whole-contig, local ±1 Mb), yielding four analysis conditions, normalising coverage to the per-species median prior to pooling so that species with different sequencing depths contribute comparably. We computed Spearman rank correlation (ρ) between each sequence feature and normalised coverage on the pooled bin set using scipy.stats.spearmanr (SciPy v1.15.2) ^97^. Bin counts per condition: IGH whole-contig 34,134 bins; IGH local 9,491 bins; IGL whole-contig 114,161 bins; IGL local 13,643 bins. Because adjacent 4 kb bins on the same contig are spatially autocorrelated, treating them as independent observations inflates the effective sample size and produces p-values well below their nominal calibration. We therefore interpret reported p-values qualitatively as evidence that ρ ≠ 0 rather than as calibrated significance levels, and treat Spearman ρ effect sizes as the primary reported quantity.

**Table S2.1:** Spearman correlation (ρ) between sequence features and normalised HiFi coverage, pooled across 24 avian species. Values computed separately for whole-contig bins and a ±1 Mb local window around the IGH or IGL locus. Note: p-values are inflated due to spatial autocorrelation between adjacent bins; effect sizes (ρ) are the relevant quantity. HP, homopolymer richness; GC, GC content; G4, G-quadruplexes; DR, direct repeats; MR, mirror repeats; APR, A-phased repeats; IR, inverted repeats; STR, short tandem repeats.

| Feature | IGH whole-contig $\rho$ | IGH local $\pm 1$ Mb $\rho$ | IGL whole-contig $\rho$ | IGL local $\pm 1$ Mb $\rho$ |
| --- | --- | --- | --- | --- |
| n (bins) | 34,134 | 9,491 | 114,161 | 13,643 |
| HP | -0.437 | -0.618 | -0.130 | -0.115 |
| GC | -0.396 | -0.312 | -0.344 | -0.239 |
| G4 | -0.034 | -0.166 | -0.183 | -0.129 |
| DR | 0.000 | -0.158 | -0.116 | -0.076 |
| MR | +0.080 | -0.146 | -0.076 | -0.066 |
| APR | +0.104 | -0.095 | +0.013 | -0.024 |
| IR | +0.208 | -0.018 | -0.087 | -0.110 |
| STR | +0.212 | -0.016 | -0.063 | -0.064 |
| Z-DNA | +0.149 | +0.065 | -0.073 | -0.053 |

**Table S2.2:**
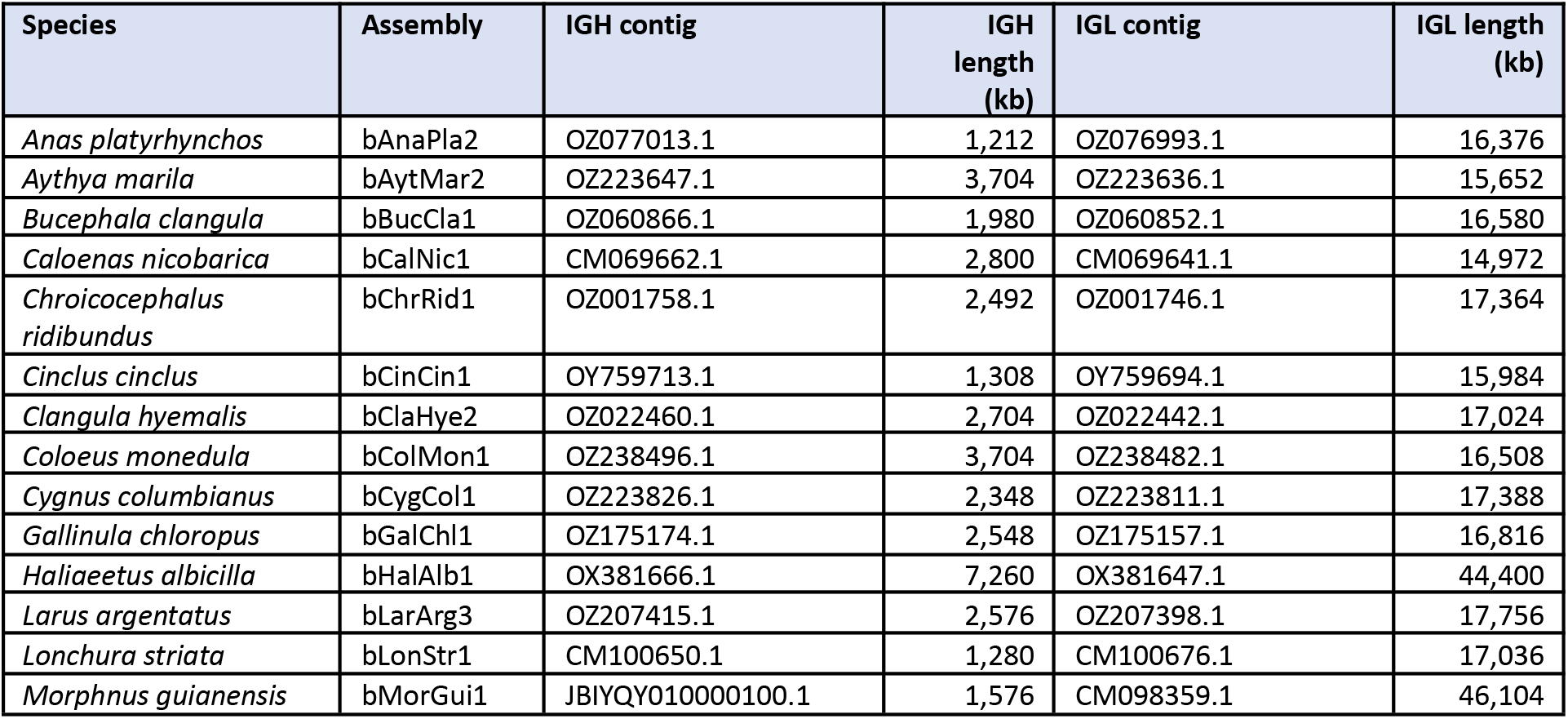
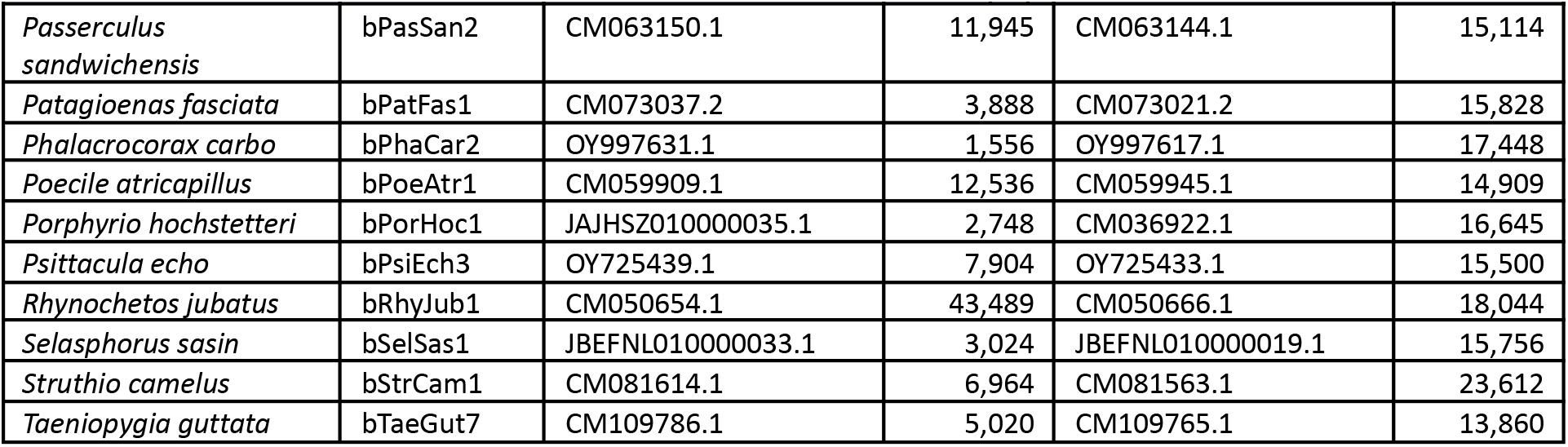
Representative contigs used for IGH and IGL coverage and sequence feature analysis.

### Supplementary Section 3: Strand distribution across the avian tree

**Figure S3:**
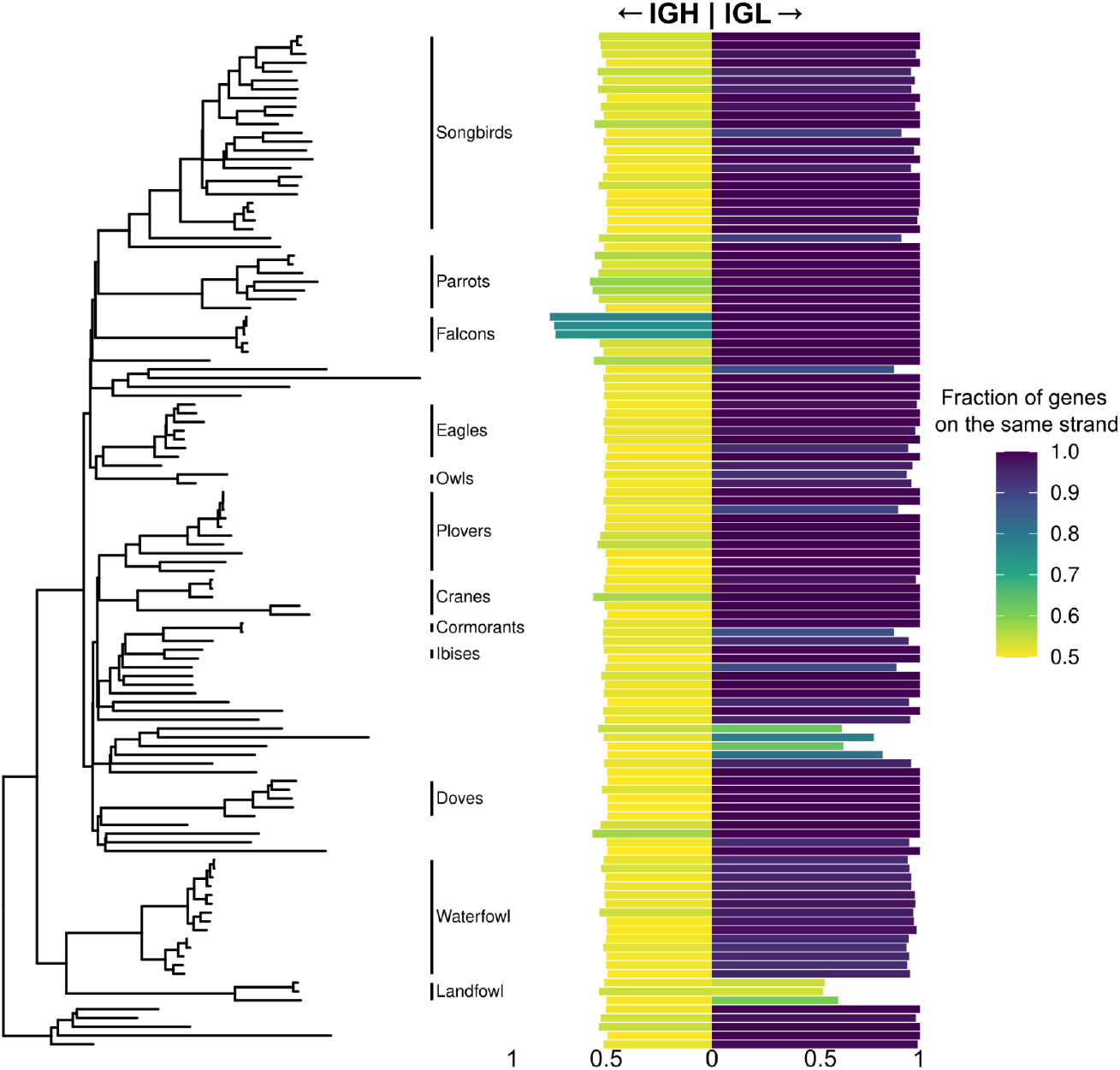
Strand distribution of genes in IGH and IGL along the bird tree. The scale goes from 0.5-1, 1 (purple) meaning all genes are located on the same strand and 0.5 (yellow) meaning half of the genes are on the positive and half on the negative strand. Orders with two or more species are indicated next to the tree.

### Supplementary Section 4: IGH locus comparison

**Figure S4:**
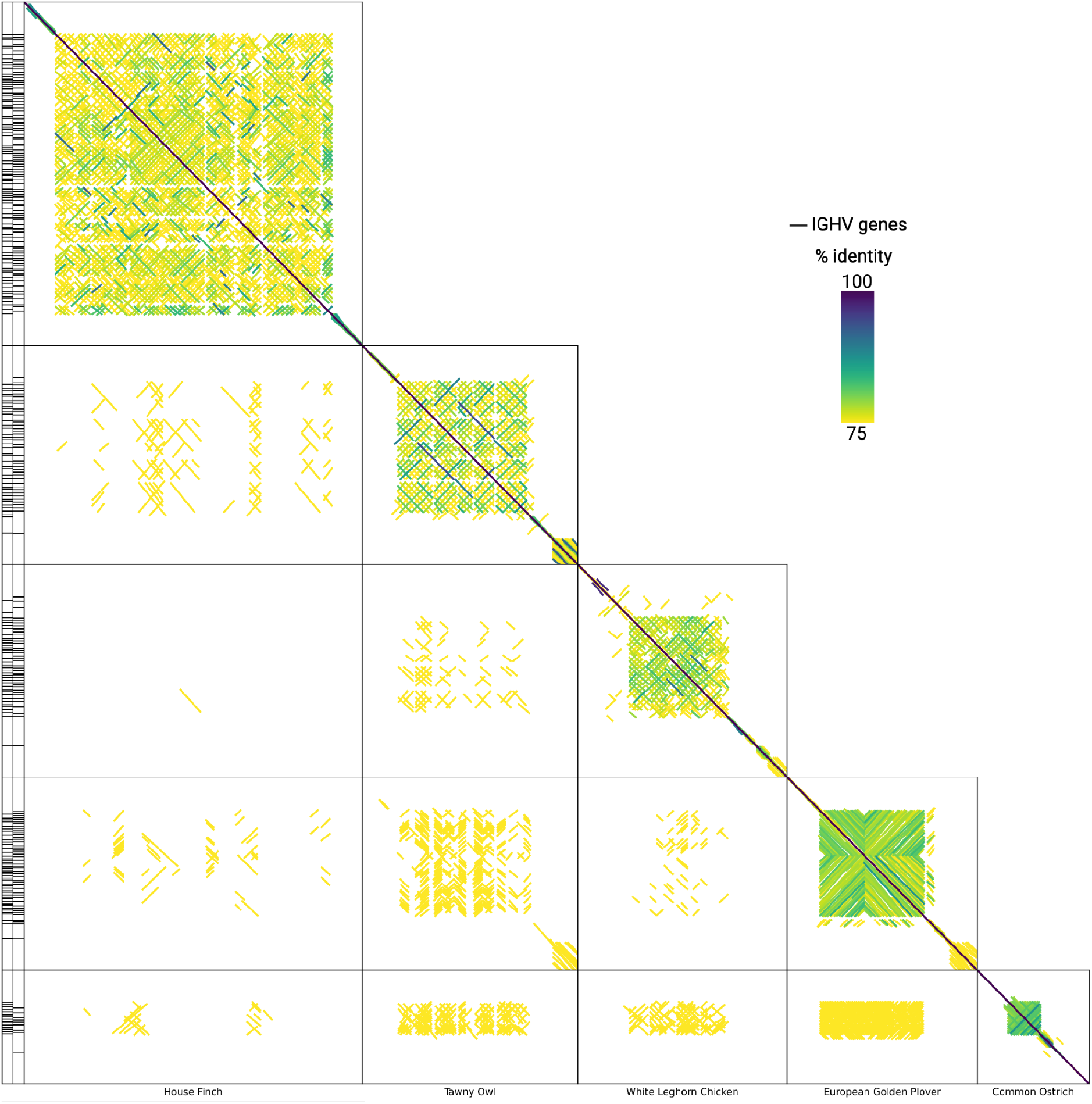
Across species comparison of IGH loci for 5 species. The dotplots show the locus similarity between different individuals.

### Supplementary Section 5: IGL locus comparison

**Figure S5.1:**
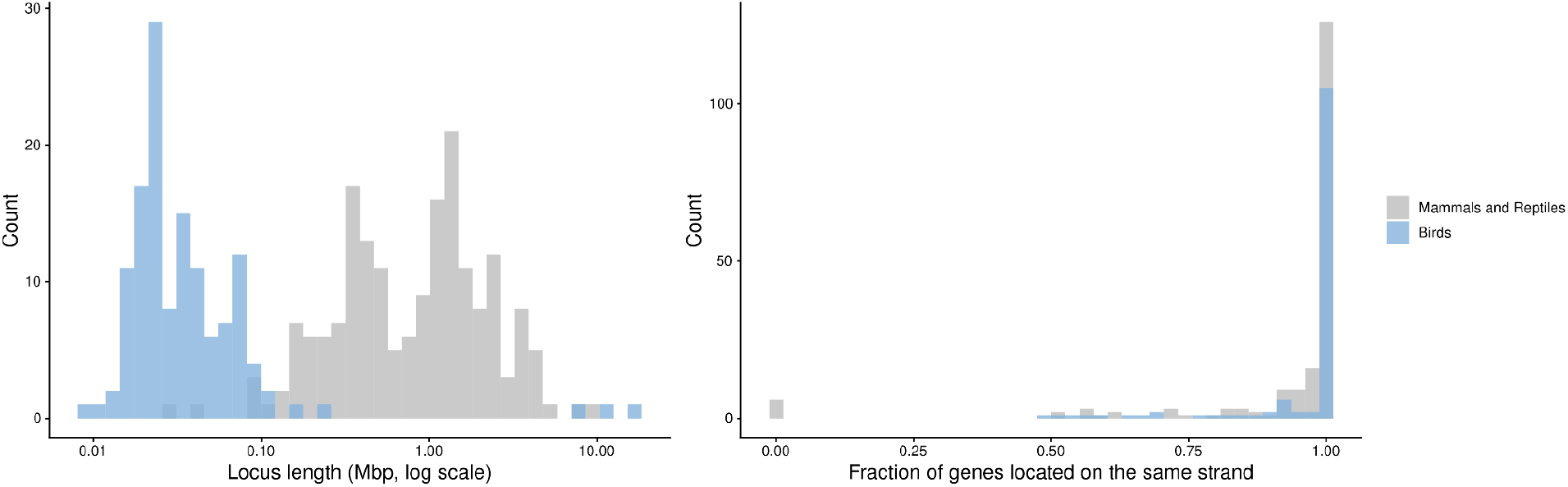
The avian IGL locus A) Locus length comparison between birds (blue) and mammals and reptiles (grey) from the VGP^3^. Length is measured in Mbp and shown on a log scale. B) Fractions of genes located on the same strand compared between birds (blue) and mammals and reptiles (grey) from the VGP. The scale goes from 0.5-1, 1 meaning all genes are located on the same strand and 0.5 meaning half of the genes are on the positive and half on the negative strand.

### Supplementary Section 6: Within-species IGH diversity (additional species)

**Figure S6.1.**
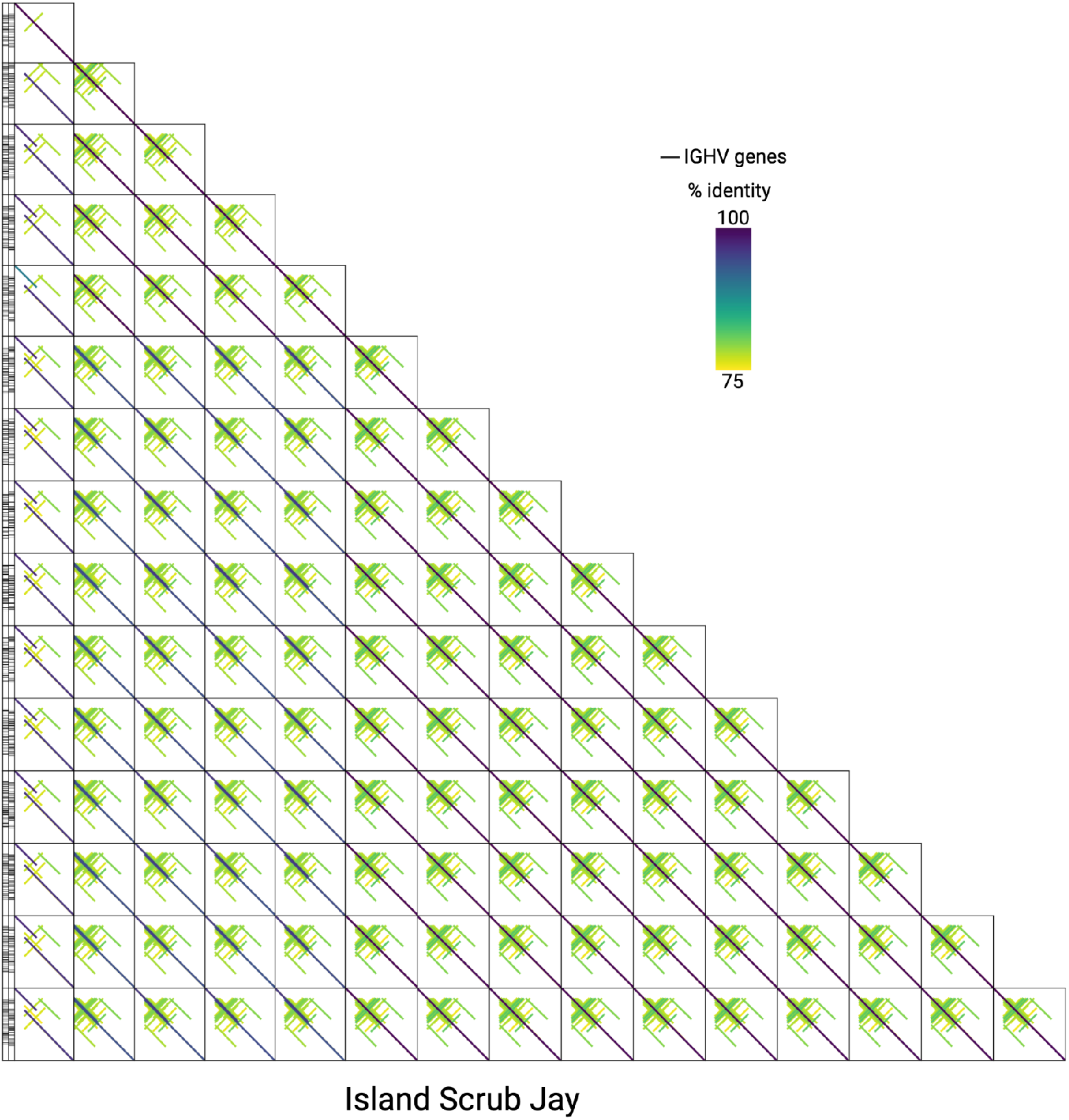
Within species comparison of IGH loci for the Island Scrub Jay. The dotplots show the locus similarity between different individuals. Only haplotypes with a confidently assembled IGH locus are shown. Haplotypes were excluded if the IGH locus was absent or if locus length and inversion content were substantially reduced relative to other haplotypes of the same species, indicative of incomplete assembly.

**Figure S6.2.**
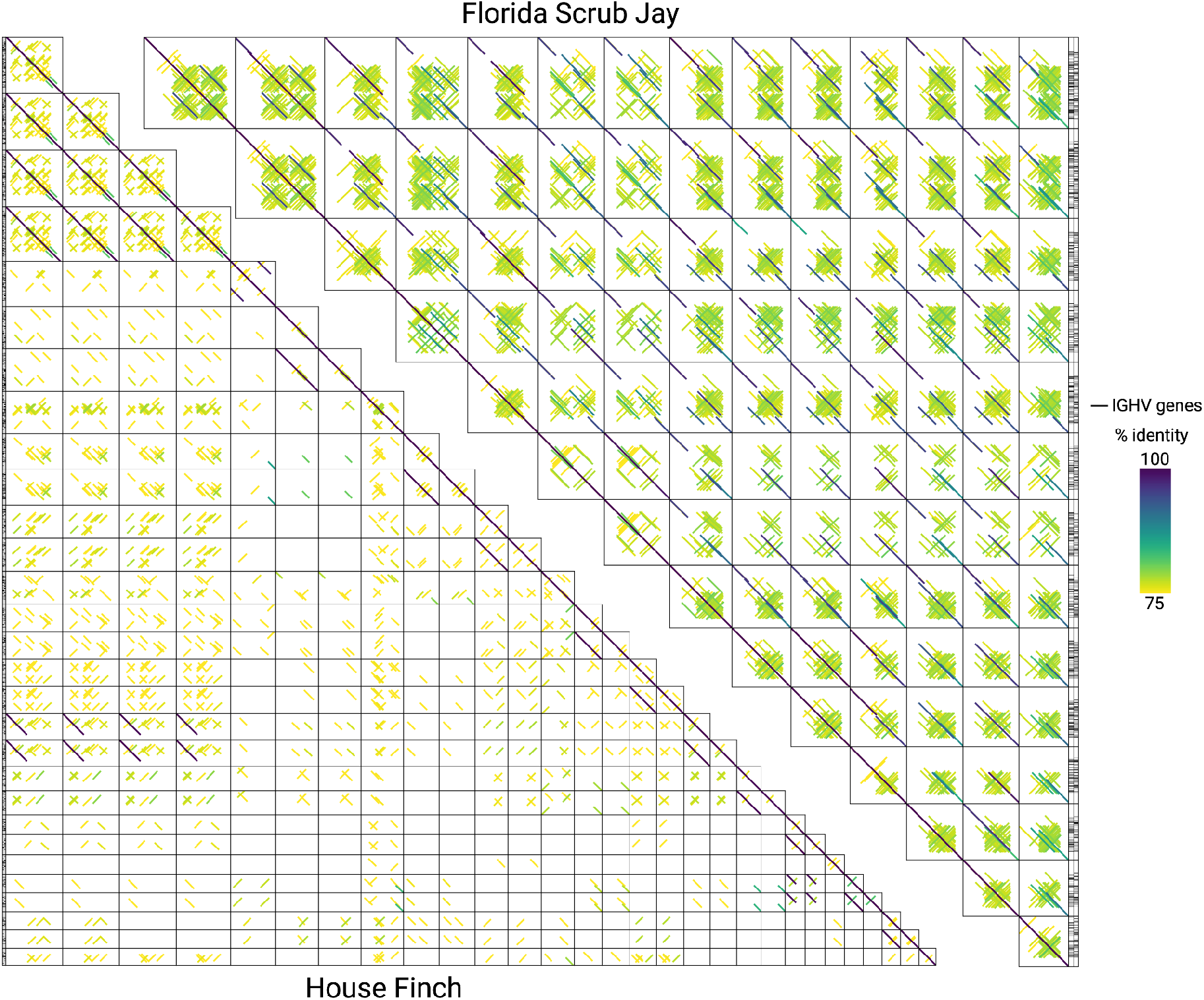
Within species comparison of IGH loci for the Florida Scrub Jay and the House Finch. The dotplots show the locus similarity between different individuals. Only haplotypes with a confidently assembled IGH locus are shown. Haplotypes were excluded if the IGH locus was absent or if locus length and inversion content were substantially reduced relative to other haplotypes of the same species, indicative of incomplete assembly.

**Figure S6.3.**
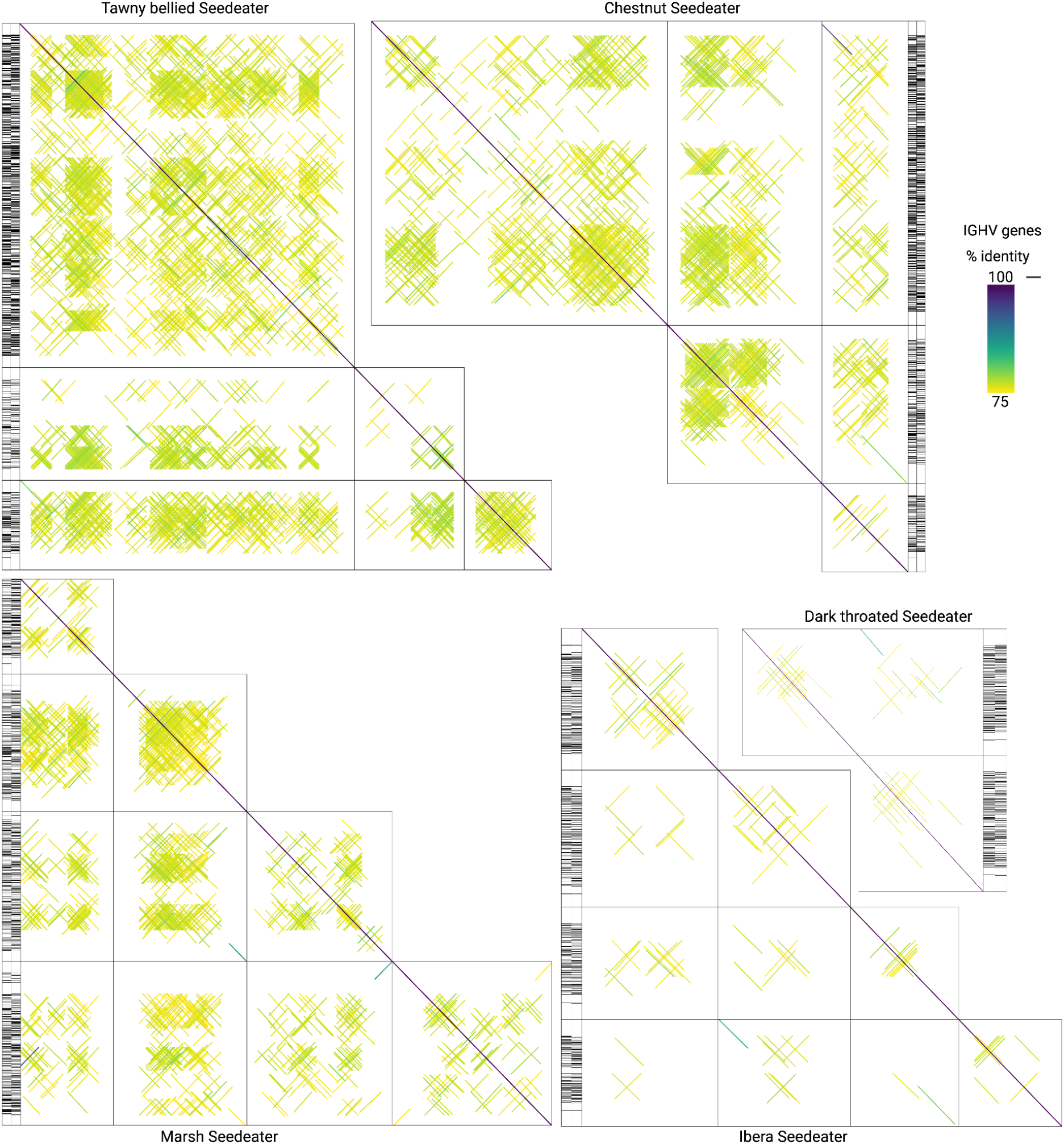
Within species comparison of IGH loci for multiple Seedeater species: Tawny bellied Seedeater, Chestnut Seedeater, Marsh Seedeater, Ibera Seedeater and Dark throated Seedeater. The dotplots show the locus similarity between different individuals. Only haplotypes with a confidently assembled IGH locus are shown. Haplotypes were excluded if the IGH locus was absent or if locus length and inversion content were substantially reduced relative to other haplotypes of the same species, indicative of incomplete assembly.

### Supplementary Section 7: Palindrome spacer analysis

**Figure S7:**
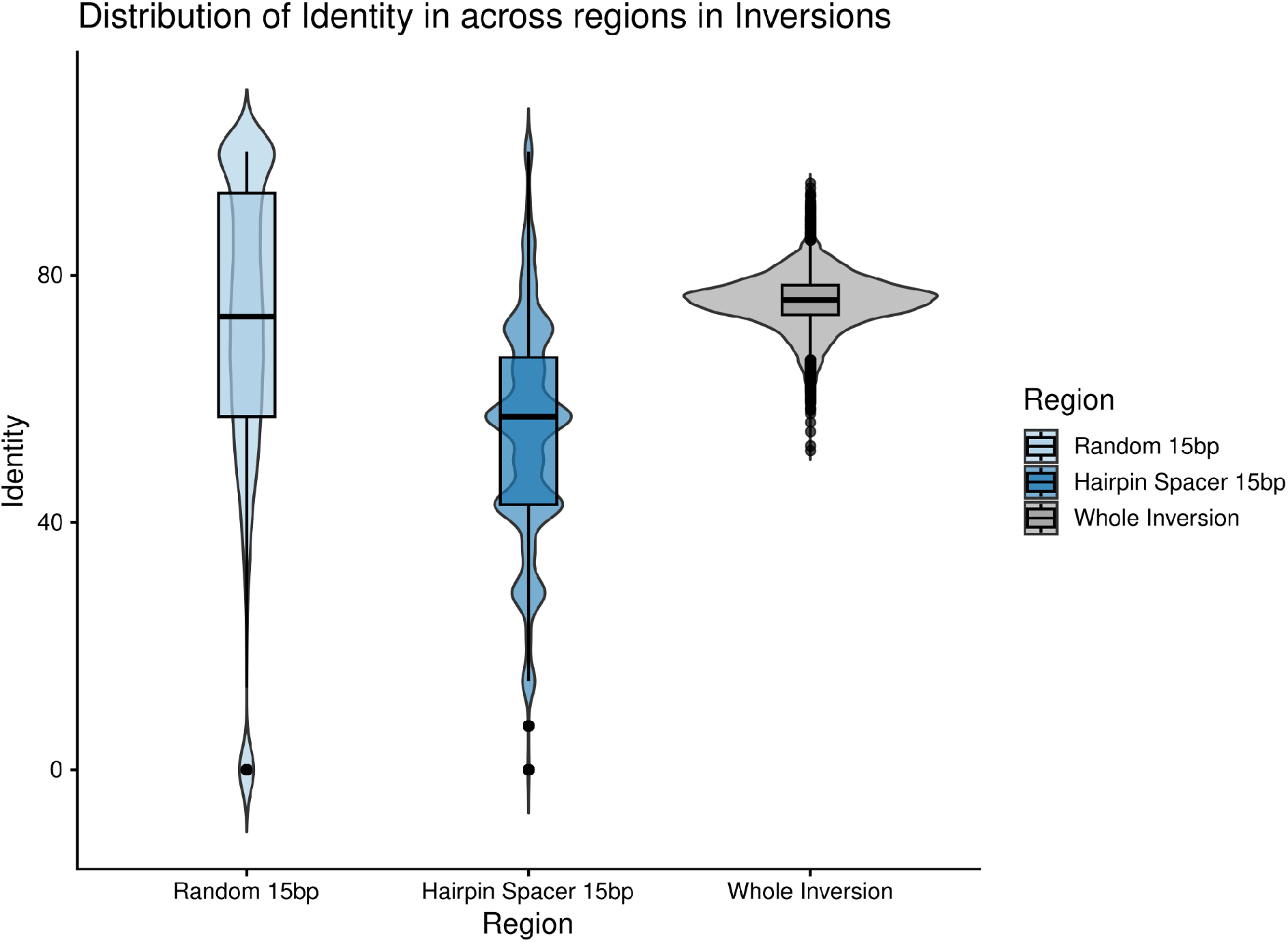
Spacer in Palindrome Sequence identity between the two arms of IGH palindromes. Every value compares a stretch of one arm with the corresponding stretch of the other: the full palindrome (grey), the central 15 bp forming the potential hairpin spacer (dark blue), and a random 15 bp window elsewhere in the same palindrome as a control (light blue).

#### Methods

For each identified inversion, we compared the palindromic identity (the sequence identity between the two halves of the inversion) across three regions: the central 15 bp, the full inversion, and a randomly selected 15 bp window. This allows us to test whether palindromic identity is reduced specifically at the center of inversions relative to both their overall sequence and to a size-matched random control.

### Supplementary Section 8: RSS motif discovery details

**Figure S8.1:**
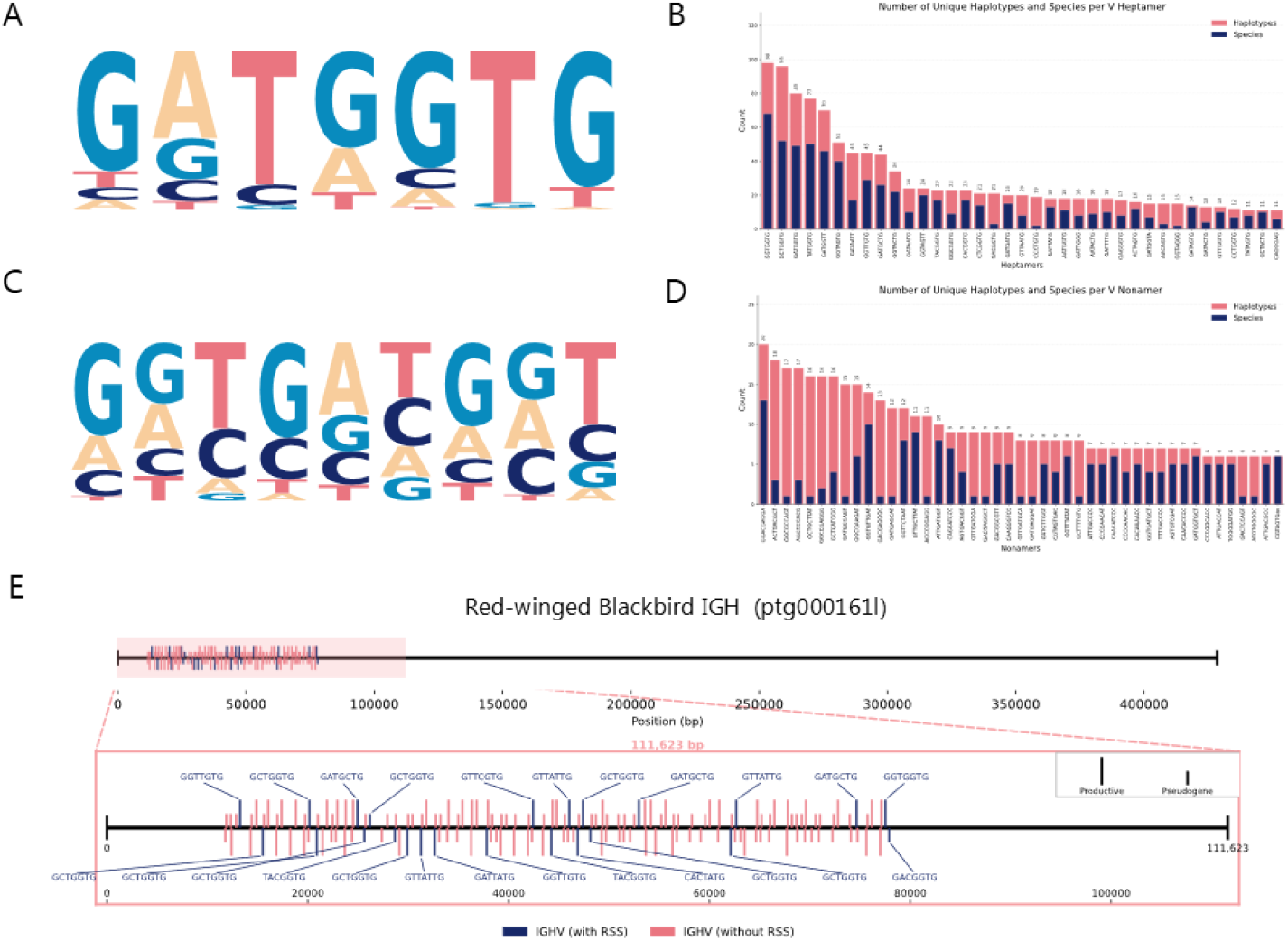
IGHV RSS analysis A) V heptamer MEME motif figure (created from heptamers appearing in more than 10 haplotypes across the dataset, n=1693). B) Bar chart showing the number of haplotypes (red) and species (blue) that top V heptamers (found in 10+ haplotypes) are found in. C) V nonamer MEME motif figure (created from nonamers appearing in more than 5 haplotypes across the dataset, n=624). D) Bar chart showing the number of haplotypes (red) and species (blue) that top V nonamers (found in 5+ haplotypes) are found in. E) Number line visualization of the annotated V cluster in the newly sequenced Red-winged Blackbird. The top horizontal line represents the cluster’s position within the contig, and the zoomed in horizontal line represents the cluster (with 10% of its size included as spacing on either side). Positive genes are shown as vertical lines above the number line, and negative genes are shown as vertical lines below the number line. Productive genes are twice the size of pseudogenes, and genes with an RSS are labeled by the heptamer present and colored blue, while genes without an RSS are unlabeled and colored red.

**Figure S8.2:**
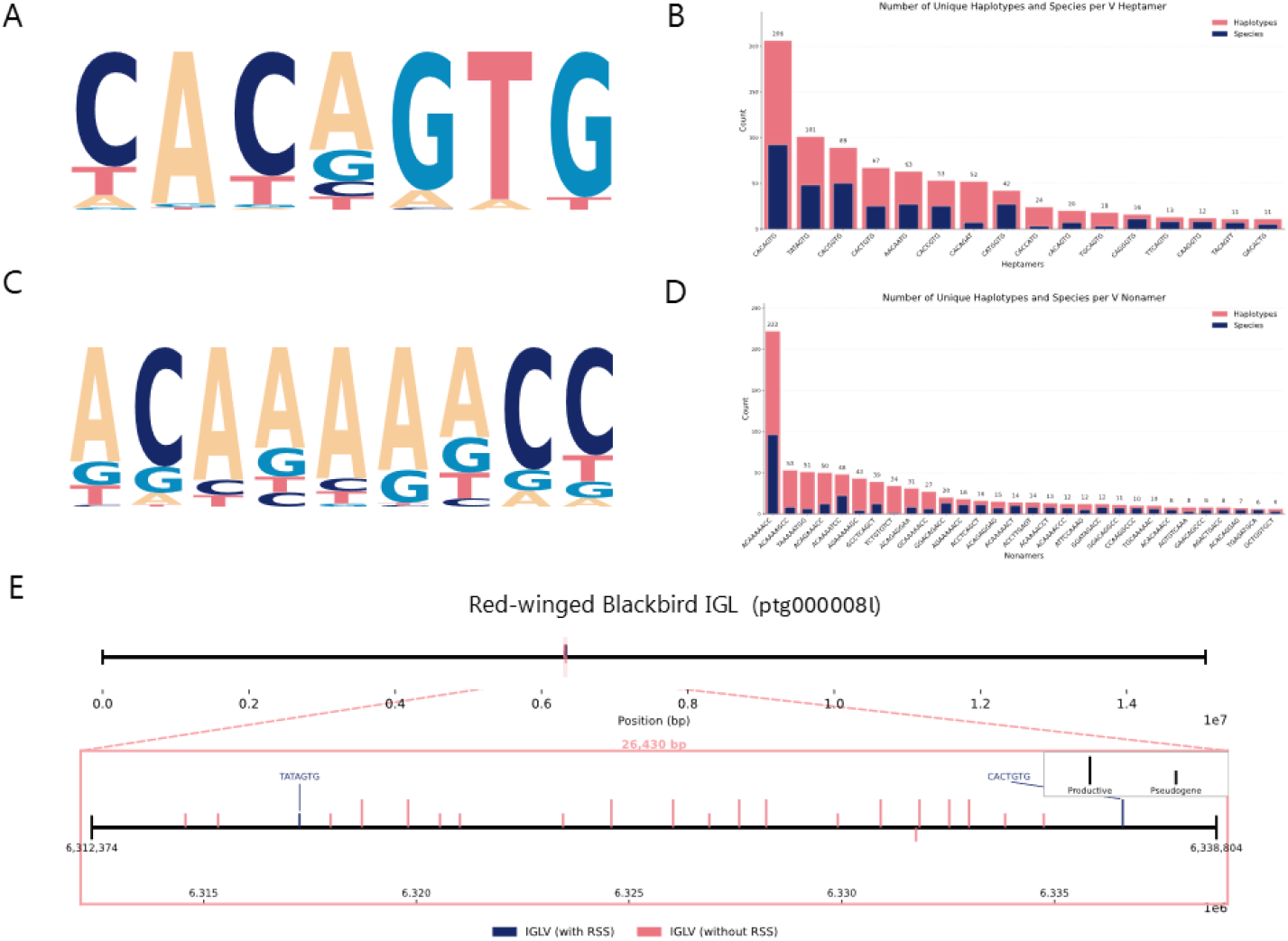
IGLV RSS Analysis Same layout as Figure S8.1 using IGLV genes, see S8.1 description for explanation (IGLV MEME heptamer n=1803, nonamer n=1625)

### Supplementary Section 9: D gene annotation and analysis

#### D Gene Annotation

While IgDetective is designed to work with all types of VDJ genes, annotation of D and J genes across the avian clade using this tool presented issues, largely due to the small size of these genes and the limited reference dataset available for them. To overcome this, we developed a separate D gene annotation procedure that searches for D RSS, as opposed to searching for actual D sequences. D genes have both upstream and downstream RSS, unlike V genes, and these follow consistent motifs, making them much easier to find than the actual D sequences^2,4^. Because of this, it is possible to find them through searching for potential upstream RSS, and then searching for a potential downstream RSS some distance away; as the sequence in between those RSS is likely to be a D gene. Using a similar RSS discovery procedure as in V genes, a D RSS reference dataset from a study on chicken IG ^69^, and distances between 20 and 70 bp, we used this method and searched for RSS in contigs where we found V genes. In cases where multiple valid downstream RSS are found off of an upstream RSS, we take only the RSS that is furthest away from the upstream RSS, as this yields the largest potential gene with the other potential valid RSS contained within it. We also applied an additional filtering step to this search, as within the reference data there was always a very high similarity between up and downstream heptamers, suggesting that potential genes with heptamers that are very different from each other likely aren’t real. We removed all potential genes with heptamers that are more than 2bp different from each other. Using a single constant heptamer and nonamer threshold did not yield the best results in all species however, so we ran the procedure using every combination of thresholds from 0-3 bp for heptamers, and 0-6 bp for nonamers. After initial testing, we decided not to include the least strict combination (3-6), as we saw a large increase in false positives at this threshold. We then analyzed the results from each combination, and took the potential genes that were found in the most strict threshold where genes were found. These genes have the highest chance of being real, as their RSS are the closest to the reference, but they often represent only a limited portion of the possible diversity of these sequences, as some real genes may have an RSS that is further away from the reference. To account for this, we also looked in every subsequent less strict combination from the first for potential genes less than 250 bp away from the initial genes (measured from the center of the gene), and took any matching this criteria. We expect D genes within a locus to be clustered together, as this is the case within the reference, so genes with more distant RSS that are very close positionally to the genes we are most confident in also have a high potential to be real.

Using this procedure yielded biologically reasonable results across the avian clade, with D genes tending to cluster very close to the V’s and the majority of them being nearly all upstream or downstream of the V’s as expected. However, many haplotypes had genes on both the positive and negative strand, which is not expected for D genes, and in many of these cases a large percentage of the opposite stranded genes were in the exact same genomic position (Figure S9.1). This is likely due to the structure of the D RSS, and the fact that many identified up and downstream RSS are nearly palindromic to each other, as shown by the main identified heptamer motifs being perfect reverse compliments of each other, and the main identified nonamer motifs being reverse compliments with a 2bp difference (Figure S9.2). This means that in many cases a potential valid RSS in the positive strand will also be a very similar RSS in the negative strand, and vice versa. Due to our procedure only considering RSS, and not the gene sequences, in these cases we cannot determine which strand is the “true” strand, so we take both. This procedure also has the potential to annotate overlapping sequences as potential genes, as it treats every base pair within a contig as the potential start to a gene, including those within already annotated sequences. We cannot reasonably determine which gene is real or functional in these cases, so we take both.

**Figure S9.1:**
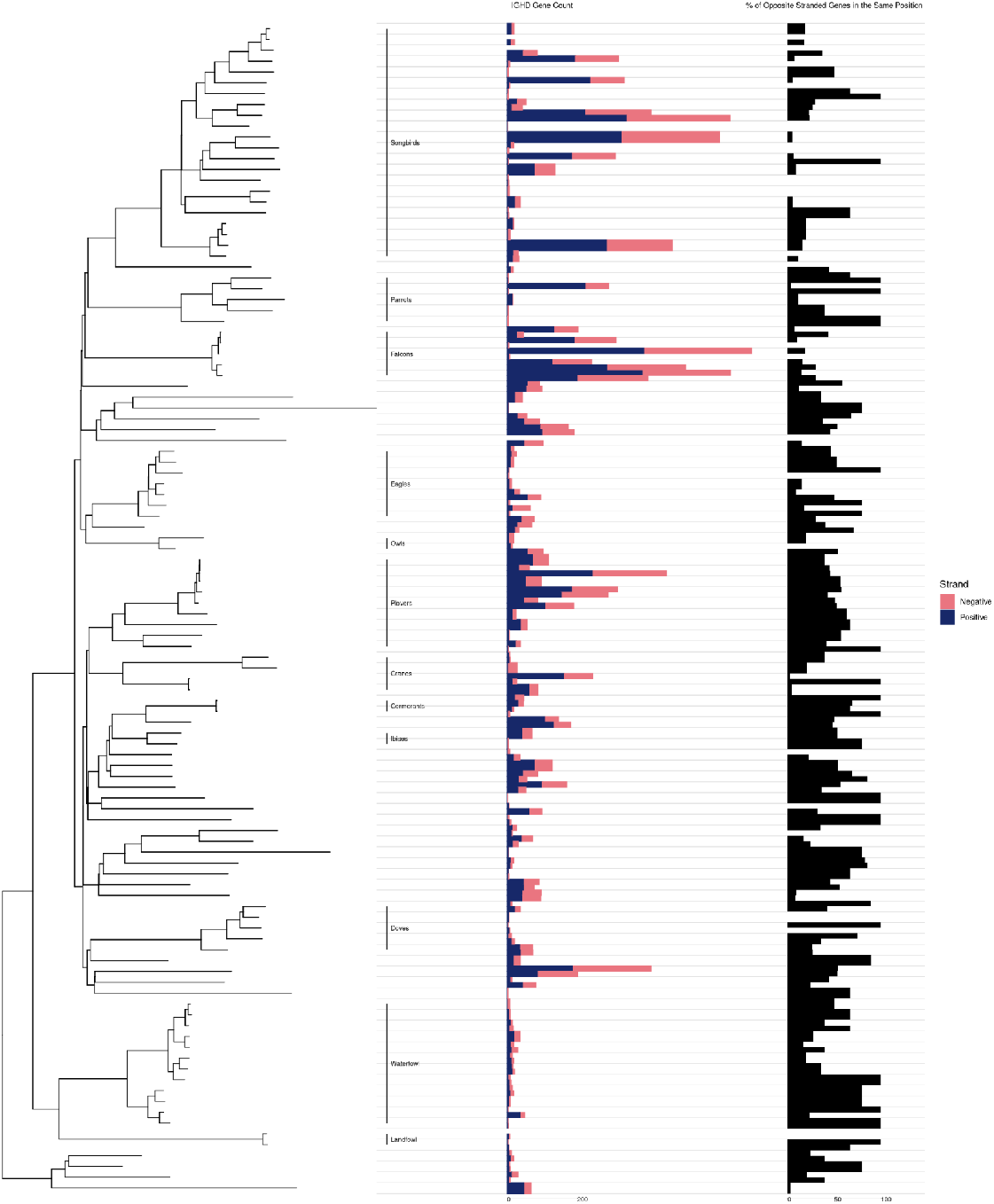
D Gene Tree IGHD haplotype level metrics from species obtained from the VGP with two or less haplotypes alongside the bird tree (labels for orders with 2 or more species provided). Light grey lines are drawn from species locations on the tree and extend through the panels, creating rows correlating to each species. If a species has both a primary and alternate haplotype, their metrics are shown in two half sized bars within that row in each panel, with the primary bar on top of the alternate. If a species has only one haplotype, one full sized bar is shown. The left column of bars represents the number of annotated V genes per haplotype, with subbars showing the number of positive (blue) and negative (red) genes. The right column of bars (black) represents the percentage of genes that are located on opposite strands but the exact same genomic position, with no bar meaning 0% of annotated genes match this criteria and a full bar meaning 100% do.

**Figure S9.2:**
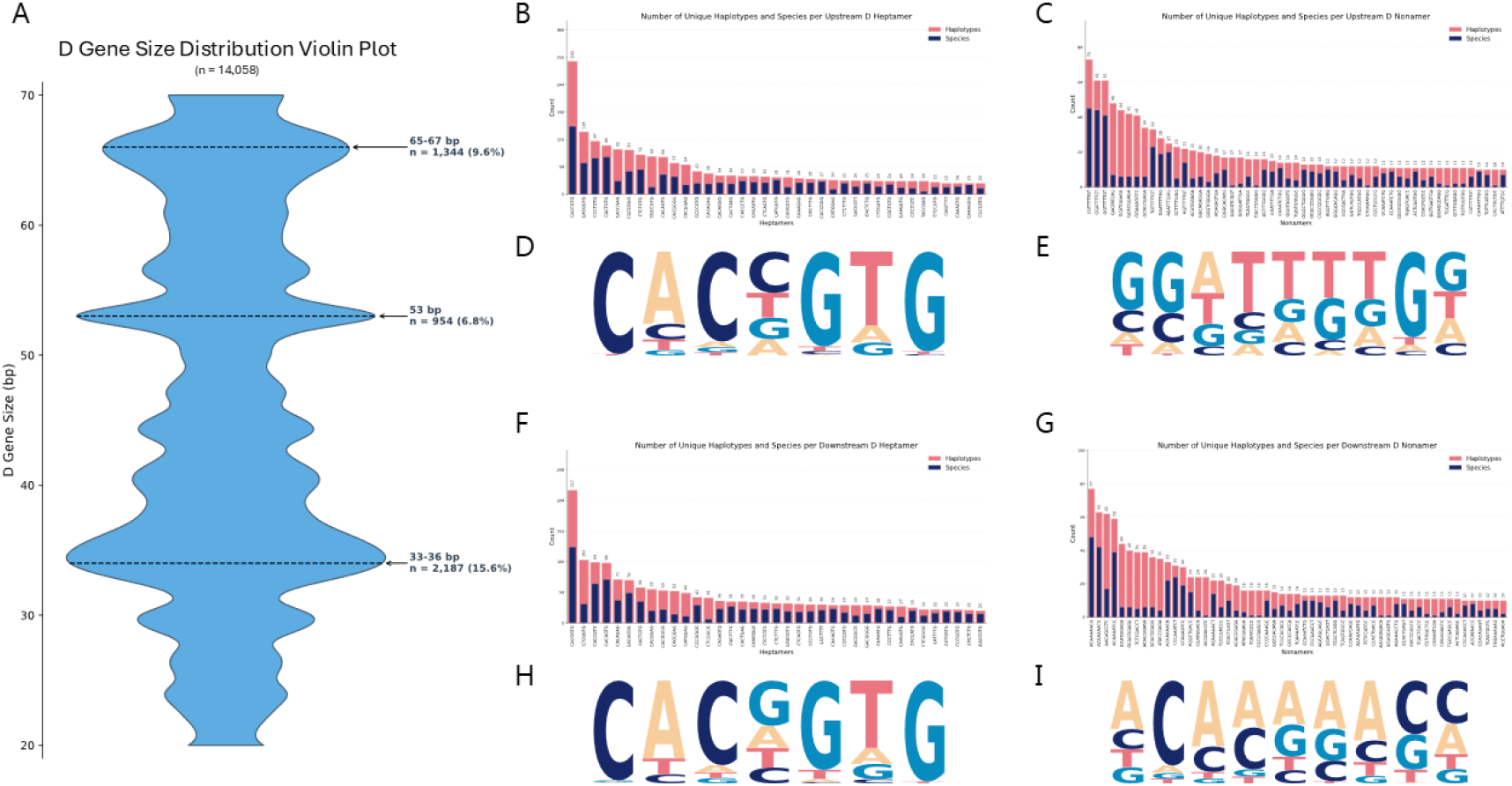
IGHD RSS and Gene Size Analysis (A) Violin plot showing the distribution of D gene sizes across the annotated dataset. The top three peaks are highlighted with a dotted black line, and labeled by the range of sizes they correlate to (bp), along with the number of genes at that size and the percentage of the dataset they make up. (B and C) Bar charts showing the number of haplotypes (red) and species (blue) that top upstream IGHD heptamers (found in 20+ haplotypes, shown in B), and upstream IGHD nonamers (found in 10+ haplotypes, shown in C) are found in. (D and E) IGHD upstream heptamer MEME motif figure (created from upstream heptamers appearing in more than 20 haplotypes across the dataset, n=8039, shown in D), and upstream nonamer MEME motif figure (created from upstream nonamers appearing in more than 10 haplotypes across the dataset, n=4496, shown in E). (F and G) Bar charts showing the number of haplotypes (red) and species (blue) that top downstream IGHD heptamers (found in 20+ haplotypes, shown in F), and downstream IGHD nonamers (found in 10+ haplotypes, shown in G) are found in. (H and I) IGHD downstream heptamer MEME motif figure (created from downstream heptamers appearing in more than 20 haplotypes across the dataset, n=8736, shown in H), and downstream nonamer MEME motif figure (created from downstream nonamers appearing in more than 10 haplotypes across the dataset, n=4555, shown in I).

### Supplementary Section 10: Red-winged Blackbird assembly quality control

Figure S10.1: Assembly quality assessment of the IGH and IGL loci in the Red-winged Blackbird (Agelaius phoeniceus, bAgePho2), generated using CloseRead^13^

**A)** Read alignment quality across the IGH locus (contig ptg000161l) for the primary haplotype. From left to right, top to bottom: mapping quality by genomic position and its frequency distribution; mismatch rate and count of indels (≥2 bp) by genomic position; counts of soft- and hard-clipped bases by genomic position; and total IGH locus length. Below, read coverage across the full locus is shown, colored by mapping quality (MapQ = 60 in grey, MapQ = 1–59 in yellow, MapQ = 0 in red), together with a basepair-resolution heatmap of local mismatch rate, in which darker shading indicates a higher percentage of mismatches per position.

**B)** As in A), for the IGL locus, shown separately for the primary (ptg000008l, blue) and alternate (atg000012l, yellow) haplotype assemblies. Locus length is compared between haplotypes using the bar chart at right.

**Figure S10.2:**
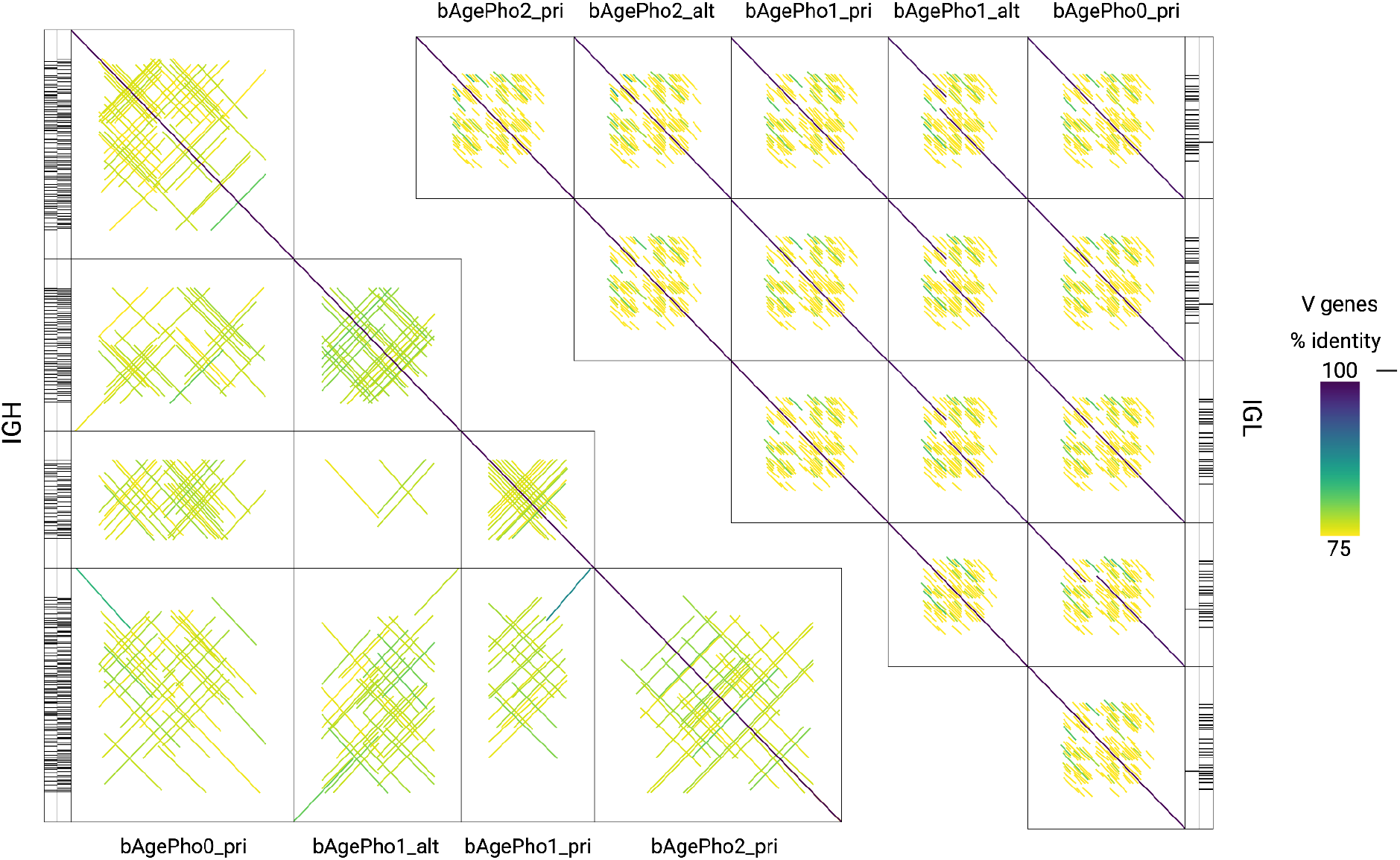
Red-Winged Blackbird Patchworkplots IGH and IGL Within species comparison of IGH loci for the Red-Winged Blackbird. The dotplots show the locus similarity between different individuals. Only haplotypes with a confidently assembled IGH locus are shown. Haplotypes were excluded if the IGH locus was absent or if locus length and inversion content were substantially reduced relative to other haplotypes of the same species, indicative of incomplete assembly.

### Supplementary Section 11: IGH donor-parent relationships

#### Correcting for clonal replication

Because a single conversion event can be represented by multiple transcripts descended from the same clonally expanded B cell, each inside-tract difference was counted once per clonal lineage per tract rather than once per transcript, to avoid pseudoreplication. Clonal relationships were inferred from the 45 bp of sequence immediately 3’ of the V gene alignment end (the region encompassing the VDJ junction), which is generated once at the point of rearrangement and inherited unchanged by all clonal descendants. Transcripts within a parent gene were greedily clustered at ≥95% junction identity to define clonal groups. This distinction was empirically necessary: in IGL, transcripts sharing a conversion tract showed junction identity indistinguishable from random transcript pairs (median 0.267 for both), indicating that shared tracts largely reflect recurrent, independent conversion events rather than clonal expansion; in IGH, by contrast, transcripts sharing a tract were substantially more clonally related (median junction identity 0.722, 50% ≥95%) than random pairs, and collapsing clonal replicates measurably changed downstream AID-spectrum results in this locus.

As an additional control against the possibility that an apparent conversion tract instead reflects the second (allelic) haplotype of the parent gene, we checked whether the donor-diagnostic positions defining each of the 20 distinct IGL tracts were present in the corresponding position of the same individual’s alternate haplotype assembly (bAgePho2_alt); none were.

**Figure S11.1:**
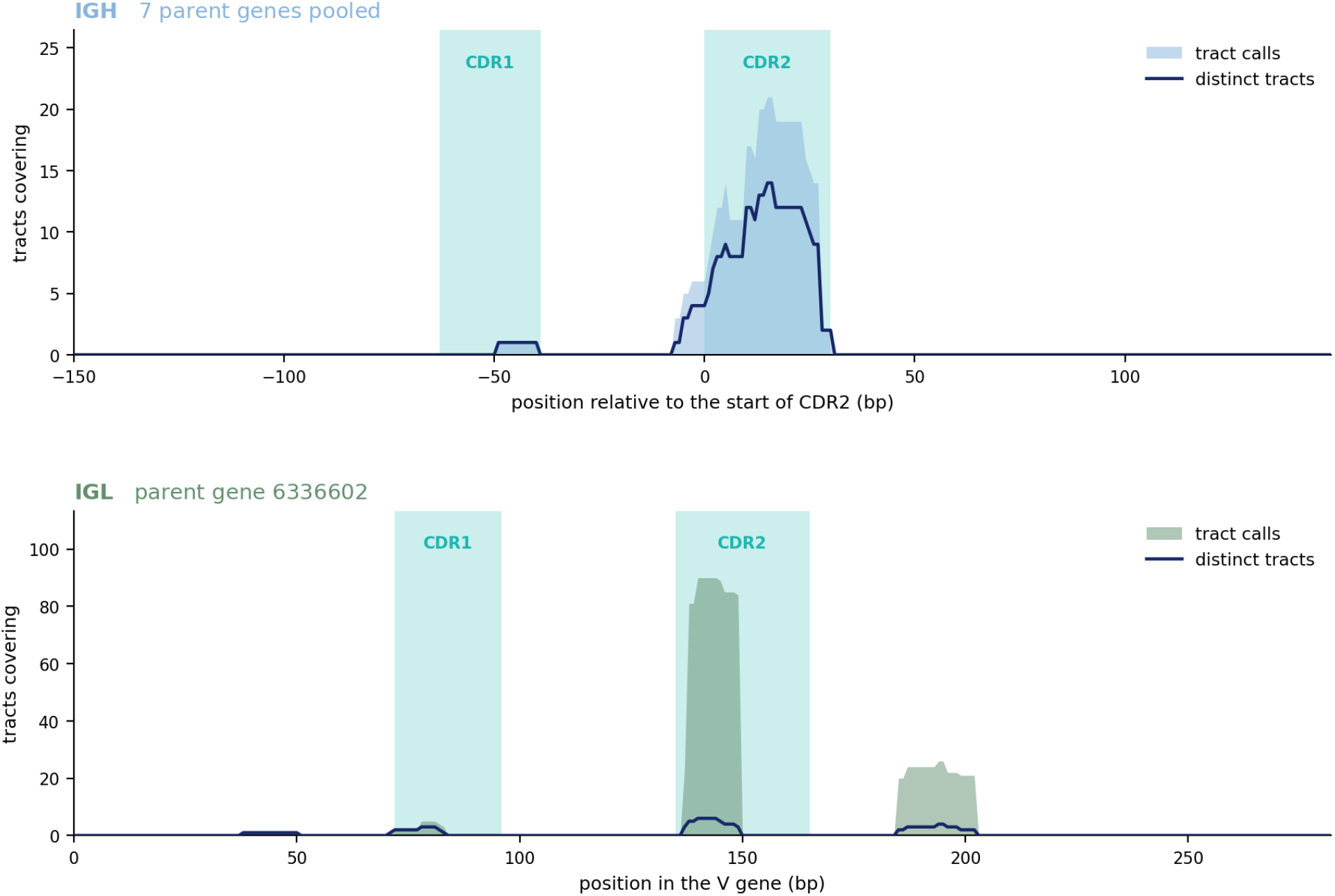
Gene conversion tracts are located in CDRs Tract calls (filled) count one per transcript-level event, using the best-supported donor where a donor is ambiguous; distinct tracts (line) count each tract position once, so clonally related transcripts collapse. Teal blocks mark CDR1 and CDR2. A) IGH. Seven parent genes pooled and drawn relative to each gene’s own CDR2 start. FR2 is 39 bp in all seven, so the CDR blocks are exact. B) IGL. the single parent gene 6336602 in its own coordinates. Only the framework boundaries are measured, from protein-level motifs^98^; CDR widths are canonical (CDR2 30 bp, CDR1 24 bp).

**Figure S11.2:**
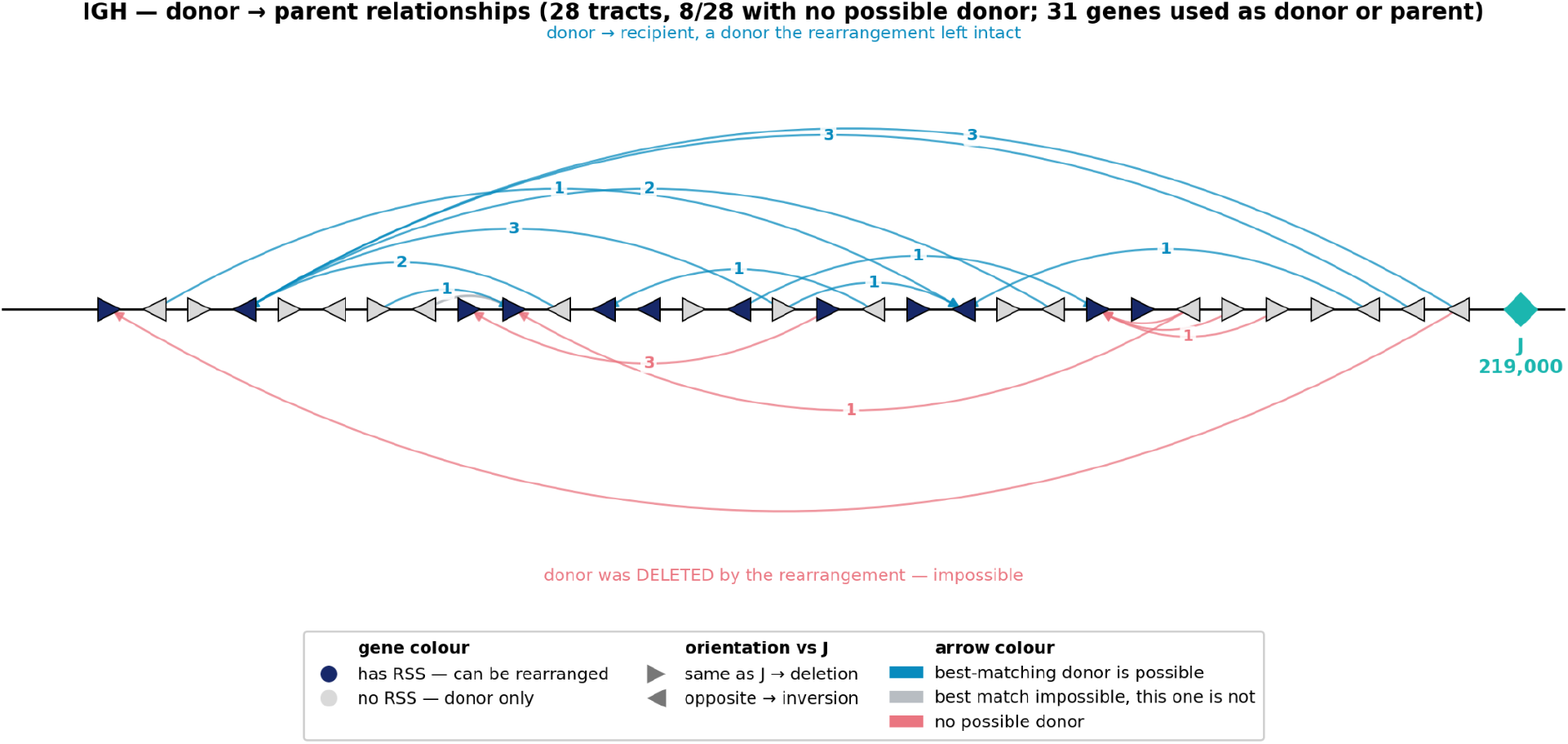
IGH donor parent relationships in the Red-winged Blackbird The 31 of 162 IGH V genes that appear as a donor or a parent, drawn at their contig positions. Fill gives RSS state (blue, RSS present and therefore rearrangeable; grey, absent — donor only); marker direction gives orientation relative to J. One arc per event, all at the same width, labelled with the number of supporting tracts: teal, the best-matching donor is one the rearrangement left intact; grey, the best-matching donor was deleted but another donor could have supplied the tract, and the arc is drawn from that surviving donor; rose, below the axis, every candidate donor had been deleted (8 of 28 events). *The J position shown (219,000) is inferred from the D cluster, not observed — no IGH J was located, and testing both J strands returns a false-discovery rate of ≈1 either way*.

#### Locating the IGH J gene

The D gene cluster was used to determine the relative position of the J gene compared to the V gene cluster.

### Supplementary Section 12: Reference-mismatch sensitivity analysis

Scoring the Red-winged Blackbird’s own transcripts against unmatched references reveals the effect of reference choice directly: when the same Red-winged Blackbird transcripts are reassigned to their best-matching V gene using germline references from other individuals rather than the matched bAgePho2 assembly, a substantial fraction are assigned to a different gene entirely (Figure S12). Even scoring against the same bird’s own second haplotype, which isolates the unavoidable cost of matching a diploid animal to a single haploid reference, produces a measurable rate of reassignment; the excess above this baseline, roughly nine percentage points in IGL, isolates the cost specifically attributable to using a different individual’s germline.

**Figure S12:**
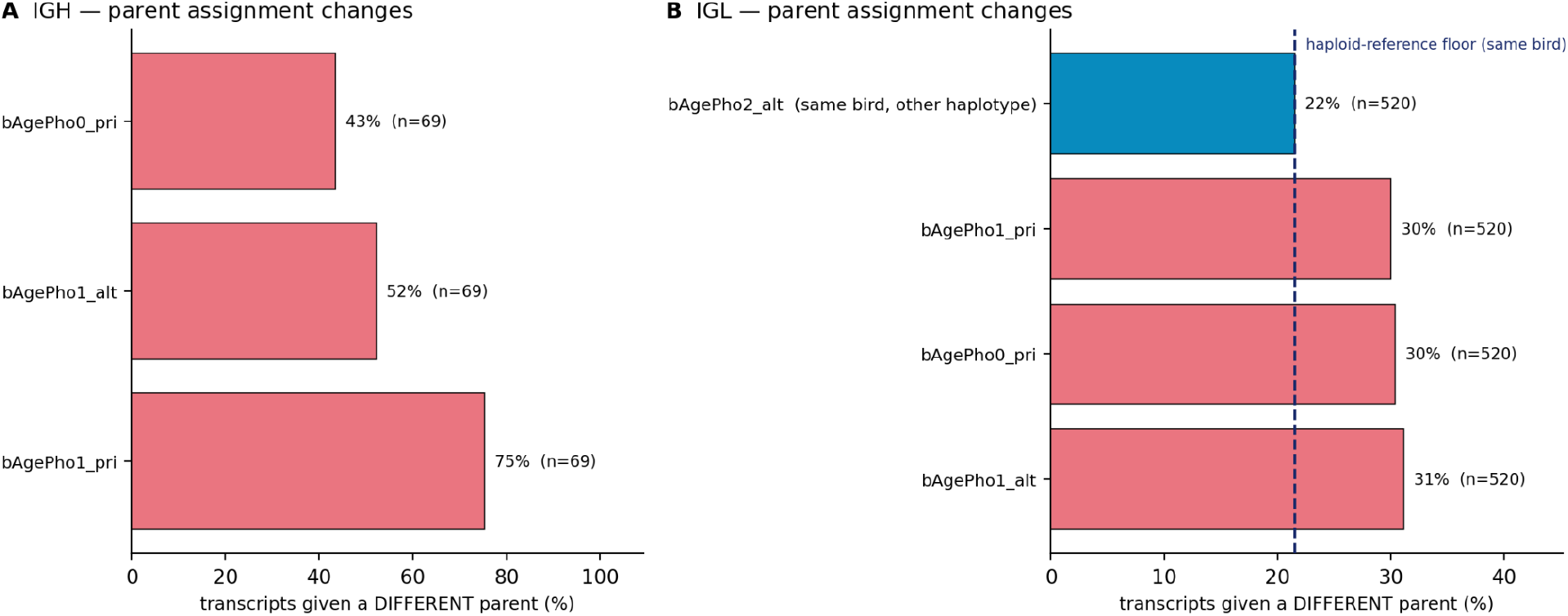
The germline reference changes which V gene a transcript is assigned to. Identical transcripts (IGH n = 69, IGL n = 520) scored against each reference and compared with the matched germline from the sequenced bird (bAgePho2_pri); orthologues paired by best reciprocal alignment. Bars give the percentage of transcripts assigned to a different V gene. (A) IGH. (B) IGL. Rose, a different individual; blue, the sequenced bird’s own second haplotype, marked by the dashed line. That line is the cost of scoring a diploid animal against a haploid reference, not a technical error floor — the matched reference pays it too. The excess above it isolates the effect of using a different animal: ∼9 percentage points in IGL. No same-bird control exists for IGH, as no IGH V genes are annotated on the alternate haplotype. Median identity to germline differs by ≤0.15 points across all references, so divergence from germline does not detect this.

